# Recombinant sialylated ApoE2 suppresses ApoE4 and sex-specifically strengthens brain metabolism and cognition in ApoE4 mice

**DOI:** 10.64898/2026.07.30.741628

**Authors:** Xin Zhang, Hee-Jung Moon, Teruna Siahaan, Liqin Zhao

**Author notes:** **Conflict of Interest Statement:** The authors have declared that no conflict of interest exists.

## Abstract

Human *APOE4* is the strongest genetic risk factor for late-onset Alzheimer’s disease (AD), whereas the relatively rare *APOE2* confers exceptional protection. This study examines whether recombinant ApoE2 protein can therapeutically leverage this genetic advantage to strengthen aging brains at risk for AD. We produced recombinant human ApoE2 (rhApoE2) using the FreeStyle 293-F system, yielding rhApoE2 with extensive sialylation that closely resembles its natural form in the human brain. In ApoE4-knockin mouse primary neurons, exposure to rhApoE2 induced significant increases in hexokinase 2 expression and decreases in endogenous ApoE4 protein levels. rhApoE2 also protected ApoE4 neurons from oligomeric amyloid-β toxicity and oxidative stress. We further developed a cadherin-derived peptide (ADTC5) that transiently increases paracellular porosity of the blood-brain barrier, enabling safe, non-invasive delivery of rhApoE2 into the mouse brain. When co-administered with ADTC5, weekly intravenous doses of rhApoE2 for 4-8 weeks in middle-aged and aged ApoE4 mice enhanced synaptosomal glycolytic and exocytotic activities, promoted cortical lipid metabolic dynamics and DHA utilization, and improved learning and memory, with sex differences observed in some outcomes. These findings offer preliminary evidence supporting the therapeutic potential of human brain-like rhApoE2 sialoprotein, which enhances metabolic robustness and cognitive strength in aging ApoE4 brains, possibly modulated by sex.

## INTRODUCTION

Alzheimer’s disease (AD) is an age-related neurodegenerative disorder characterized in part by progressive, region-specific brain glucose hypometabolism that precedes the onset of clinical symptoms [1, 2]. Impaired glycolysis leads to elevated brain glucose levels that correlate with the severity of amyloid plaque deposition and neurofibrillary tangle formation in AD patients [1]. By contrast, enhanced glycolysis was found to confer neuroprotection against amyloid toxicity and promote memory acquisition [3–5]. These findings establish glycolysis as a critical player in maintaining brain glucose homeostasis and a key determinant of brain resilience against AD.

*APOE* is currently considered the most significant genetic risk factor for AD development, with ApoE4 conferring the highest risk, whereas ApoE2 is clinically proven to be robustly protective [6–8]. One major risk associated with ApoE4 is impaired cerebral glucose metabolism during aging, independent of Aβ pathology [9]. Individuals with mild cognitive impairment (MCI) who carry the ApoE4 allele show further reductions in the cerebral metabolic rate of glucose (CMRglc) compared with non-ApoE4 carriers, and ApoE4-associated cerebral hypometabolism is linked to subjective memory decline [10, 11]. Intriguingly, cognitively intact ApoE4 carriers exhibit cerebral glucose metabolic patterns resembling those observed in clinically symptomatic AD patients, suggesting that glucose metabolism may be a major driver of ApoE-related risk for AD [12, 13]. These clinical observations highlight a regulatory role for ApoE in brain glucose metabolism; however, the underlying mechanisms and how ApoE isoforms differ remain unclear. We previously demonstrated that ApoE2-expressing mouse brains exhibited the most robust glycolytic profile compared with ApoE3- and ApoE4-expressing mouse brains, as evidenced by increased hexokinase (HK) protein expression and enzymatic activity, enhanced glycolytic flux, and elevated ATP production [14]. We further showed that ApoE2 overexpression, but not ApoE3, rescued ApoE4-induced glycolytic deficits [15], yet the therapeutic role of ApoE2 remains an open question. The present study sought to further evaluate whether ApoE2 can be therapeutically leveraged to restore glycolytic metabolism and neuronal resilience in ApoE4-expressing neuronal cultures and mouse models.

To achieve this goal and establish human relevance, we first produced physiologically relevant recombinant human ApoE2 (rhApoE2) with post-translational modifications (PTMs) that resemble those of endogenous ApoE2 in the human brain. Human ApoE proteins differ from murine ApoE at multiple levels, with one major difference being that murine ApoE undergoes minimal glycosylation, whereas human ApoE is highly glycosylated [16]. We recently reported that in the human brain, ApoE2 exhibits the most abundant sialylation, whereas ApoE4 is the least sialylated, and that the sialic acid moiety on human ApoE modulates ApoE interactions with Aβ peptides and Aβ fibrillation [17]. These novel findings provide a compelling explanation for the well-documented differential roles of human ApoE isoforms in Aβ pathogenesis in AD. Regrettably, commercially available rhApoE2 produced primarily in *E. coli* lacks the machinery for human protein PTMs, limiting the translational relevance of prior studies. We therefore established an expression system by constructing a mammalian expression vector for ApoE2 with the C-terminus fused to a Strep-tag, pcDNA3.1(-)-hApoE2-Strep. FreeStyle 293-F cells, a suspension culture expression system, were transfected to express secreted rhApoE2 in the culture medium, followed by purification using a gravity-flow Strep-Tactin Superflow column. PTM characterization confirms that rhApoE2 derived from the 293-F system is extensively modified by *O*-linked glycosylation and sialylation, which mimic the physiological profiles of ApoE2 expressed in the human cortex [17].

Here, we report the first investigation of the therapeutic potential of the human brain-like rhApoE2 sialoprotein in ApoE4-expressing mouse models. The findings reveal that rhApoE2 boosts neuronal hexokinase and promotes neuronal survival against neurotoxic insults. Importantly, it also reduces endogenous ApoE4 levels in primary neurons expressing ApoE4. When administered intravenously with the BBB modulator ADTC5, rhApoE2 induced neural responses suggesting enhanced brain glycolytic activity, synaptic vesicle release, lipid metabolic adaptability, and DHA utilization, which may together contribute to cognitive improvements. Some effects showed sex-specific differences, aligning with growing evidence of the complex interplay among sex, ApoE genotype, and AD risk.

## RESULTS

### rhApoE2 upregulated HK2 expression and Akt phosphorylation and downregulated endogenous ApoE4 levels in ApoE4-expressing primary neurons

In the brain, ApoE is primarily synthesized by astrocytes, but it is also produced in neurons, especially under stress, injury, and with aging. Recent studies have identified neuronal ApoE as a critical regulator of neuronal vulnerability in AD, and neuron-specific ApoE4 knockout was shown to mitigate neuronal, synaptic, and hippocampal volume loss in aged ApoE-KI mice [18, 19]. Moreover, neuronal glycolysis is differentially regulated by ApoE isoforms, with ApoE2 enhancing and ApoE4 suppressing glycolysis, thereby exerting a significant influence on neuronal health during aging [14, 15]. To investigate whether rhApoE2 can counteract ApoE4-mediated glycolytic deficits and improve neuronal health, we examined glycolytic changes in response to rhApoE2 treatment in primary cortical neurons derived from hApoE4KI mice. As described above, rhApoE2 was produced using the FreeStyle 293-F mammalian cell expression system. The sialylation profile of the resulting rhApoE2 protein was confirmed by neuraminidase treatment, which caused the disappearance of the upper sialylated bands (Fig. 1A). Neurons were isolated from postnatal day 0-2 hApoE4KI mouse brains (Fig. S1A-S1D) and cultured for up to 25 days in vitro (DIV). We measured glycolytic markers in these cells and observed a time-dependent increase that peaked at 23 DIV (Fig. S1E-S1F). To investigate the impact of rhApoE2 on ApoE4-expressing neurons, hApoE4KI primary neurons were treated with 25 μg/ml or 100 μg/ml rhApoE2 for 2 days (Fig. 1B). Cell lysates were collected and analyzed for protein expression. The results showed a significant dose-dependent increase in ApoE and HK2 expression with rhApoE2 treatment, while HK1 remained unchanged. Notably, endogenous ApoE4 expression decreased markedly in a dose-dependent manner (Fig. 1C-1E). Additionally, the PI3K/Akt signaling pathway was upregulated, as indicated by an increased pAkt (Ser473)/total Akt ratio (Fig. 1F). These findings align with our previous study and demonstrate that rhApoE2 treatment can induce HK2 protein expression in hApoE4-expressing neurons, potentially through activation of the PI3K/Akt signaling pathway [15]. To further confirm the impact of ApoE2 on endogenous ApoE4 expression, N2a cells stably expressing human ApoE4 were transfected with a mammalian expression vector encoding human ApoE2 (group: ApoE4 + ApoE2) or an empty vector (group: ApoE4 + V). Intriguingly, we observed a dose-dependent decrease in endogenous ApoE4 protein levels as ApoE2 expression increased (Fig. 1G-1I). Collectively, these data demonstrate that rhApoE2 can downregulate endogenous ApoE4 and rescue ApoE4-associated deficits in HK2 expression, potentially through PI3K/Akt signaling.

**Fig. 1.**
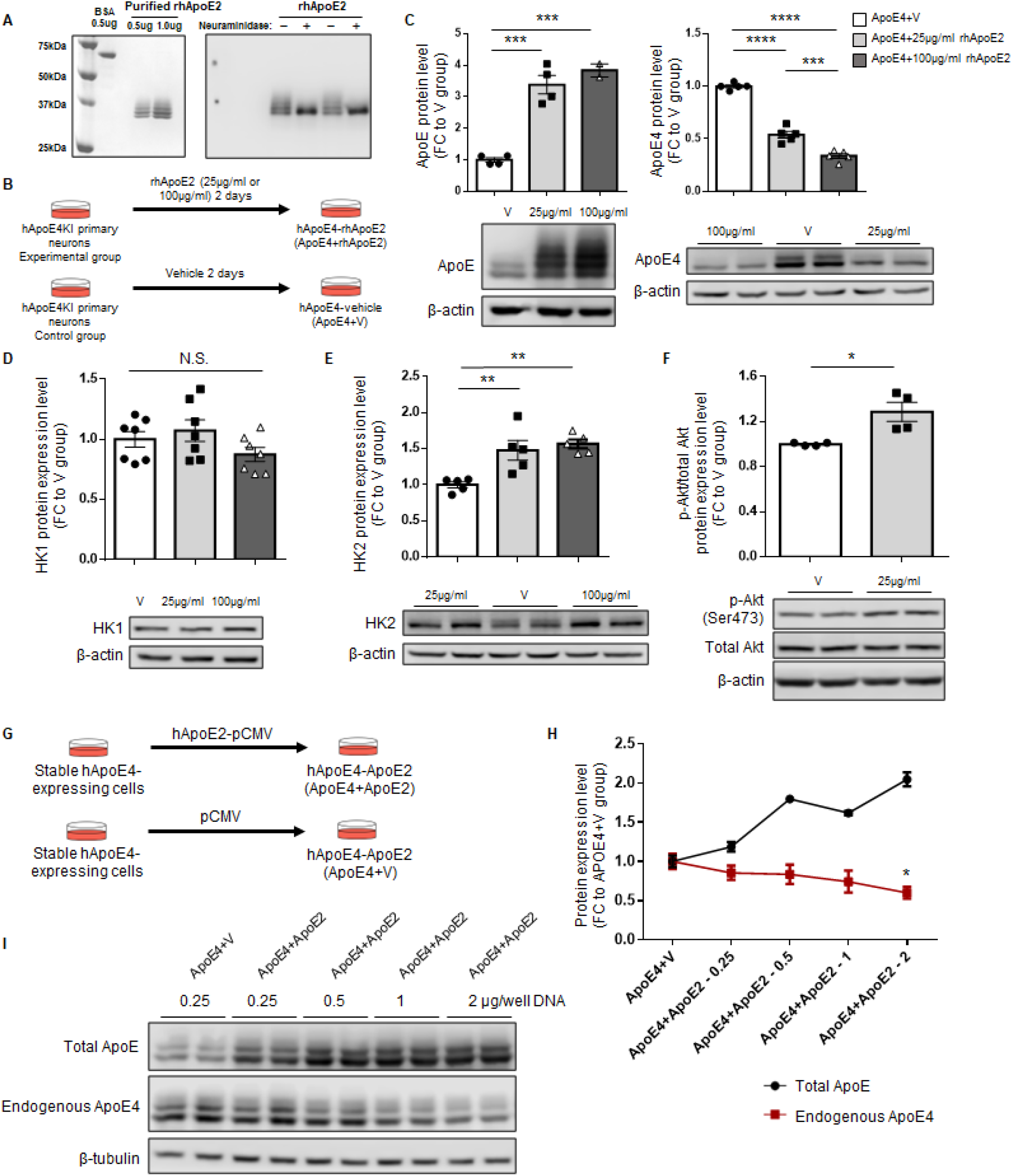
rhApoE2 upregulated HK2 expression and Akt phosphorylation while downregulating endogenous ApoE4 levels in ApoE4-expressing neuronal models. (**A**) The purity and sialylation status of human-comparable rhApoE2 were assessed by SDS-PAGE using the InstantBlue Coomassie protein stain. Pure BSA served as the reference standard. Sialylation of rhApoE2 was evaluated by neuraminidase treatment and confirmed by immunoblotting using anti-ApoE antibodies. (**B**) Schematic experimental design for rhApoE2 treatment in hApoE4KI primary neuronal cultures under normal conditions. (**C-F**) Western blot data showed that 2 days of rhApoE2 treatment led to significant increases in total ApoE, HK2 expression, and Akt phosphorylation, and to drastic decreases in endogenous ApoE4 levels in hApoE4KI primary neurons. (**G**) Illustration of N2a-hApoE4 stable cells transfected with mammalian expression vectors encoding human ApoE2 (ApoE4+ApoE2) or an empty vector (ApoE4+V). 48–72 hr after transfection, cells were analyzed for total ApoE and endogenous ApoE4 protein expression. (**H-I**) Western blot data showed that endogenous ApoE4 levels gradually decreased as ApoE2 expression increased in N2a-hApoE4 stable cells. *p<0.05, **p<0.01, ***p<0.001, ****p<0.0001. Data are presented as the group mean ± SEM. Group comparisons were analyzed using one-way ANOVA with Tukey’s post hoc or Student’s t-test.

### rhApoE2 promoted neuronal survival and HK2 expression and suppressed endogenous ApoE4 increases in response to neurotoxic insults in ApoE4-expressing primary neurons

The positive impact of rhApoE2 was further validated by its ability to confer neuroprotection to ApoE4-expressing primary neurons against neurodegenerative insults, including H_2_O_2_ and oligomeric amyloid β_1-42_ (oAβ; Fig. 2). hApoE4KI primary neurons were pretreated with rhApoE2 or vehicle for 2 days, then exposed to 100 μM H_2_O_2_ or vehicle for 30 min and finally replaced with rhApoE2 or vehicle for 6 hr (Fig. 2A). Results showed that rhApoE2 treatment significantly reduced H_2_O_2_-induced neuronal cell death, as evidenced by decreased LDH release and improved cellular metabolic activity (Fig. 2B-2C). Western blot analysis of cell lysates showed increased NeuN levels, indicating improved neuronal survival (Fig. 2D). Similar results were observed in neurons exposed to oAβ (Fig. 2E-2H and Fig. S2). Notably, endogenous ApoE4 levels increased in a dose-dependent manner in response to oAβ, consistent with previous studies showing that ApoE synthesis is induced in neurons under stress conditions (Fig. 2I-2K) [20].

**Fig. 2.**
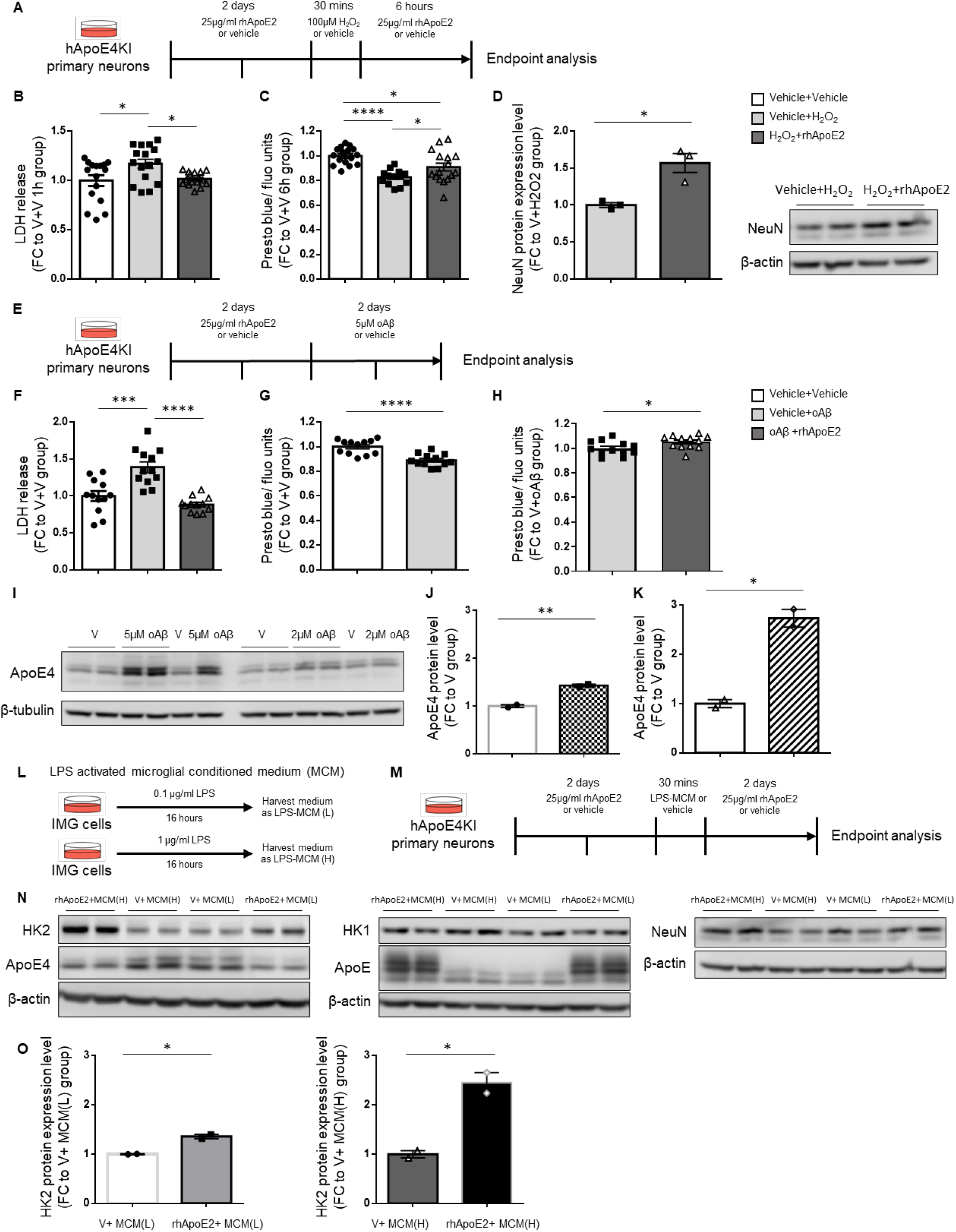
rhApoE2 promoted neuronal survival and HK2 expression and suppressed endogenous ApoE4 increases in response to neurotoxic insults in primary neurons derived from hApoE4KI mice. (**A**). Schematic experimental design for investigating the neuroprotective effects of rhApoE2 on hApoE4KI primary neurons under H_2_O_2_ challenge. (**B-D**) rhApoE2 increased neuronal viability against H_2_O_2_-induced neurotoxicity, as measured by LDH release as a marker of neuronal membrane integrity, Prestoblue fluorescence as an indicator of neuronal metabolic activity, and NeuN expression. (**E**) Schematic experimental design for investigating the neuroprotective effects of rhApoE2 on hApoE4KI primary neurons under oligomeric Aβ (oAβ) challenge. (**F-H**) rhApoE2 increased neuronal viability against oAβ-induced neurotoxicity. (**I-K**) Endogenous ApoE4 increased in a dose-dependent manner in response to oAβ. (**L-M**) Illustration of LPS-activated microglial conditional medium (LPS-MCM) generation and the experimental design to assess the effects of rhApoE2 on hApoE4KI primary neurons in response to LPS-MCM. (**N-O**) rhApoE2 increased HK2 expression and suppressed ApoE4 increases in hApoE4KI neurons under LPS-MCM challenges. \**p*<0.05, \*\**p*<0.01, \*\*\**p*<0.001, \*\*\*\**p*<0.0001. Data are presented as the group mean ± SEM. Group comparisons were analyzed using one-way ANOVA with Tukey’s post hoc or Student’s t-test.

Microglial overactivation is a well-known risk factor for neuronal apoptosis and for exacerbating AD pathology [21]. Immortalized microglial (IMG) cells were activated with 0.1 μg/ml (low dose) or 1 μg/ml (high dose) lipopolysaccharide (LPS) to evaluate the effects of rhApoE2 on primary neurons expressing ApoE4 in response to microglial inflammation. LPS-activated microglia-conditioned medium (LPS-MCM) was collected, diluted in primary neuron culture medium, and used to challenge ApoE4-expressing neurons, with or without rhApoE2 (Fig. 2L-2M). There appeared to be a synergistic effect of rhApoE2 and LPS-MCM on HK2 modulation, with high-dose LPS-MCM augmenting the rhApoE2-mediated upregulation of HK2 more than low-dose conditioned medium did (Fig. 2N-2O). The NeuN level was significantly increased, indicating that rhApoE2 protected against LPS-MCM-induced neuronal loss (Fig. 2N). Overall, the data support a neuroprotective role for exogenous rhApoE2 treatment against neurodegenerative insults in primary neurons expressing ApoE4, including oxidative stress and amyloid-associated neurotoxicity.

### Blood-brain barrier modulating peptide ADTC5 improved brain delivery of intravenously administered rhApoE2 in adult hApoE4KI mice

A major challenge in the present study was delivering rhApoE2 to the brain, as the blood-brain barrier (BBB) restricts the entry of most peripheral molecules. One of the most successful and non-invasive clinical methods for drug delivery is osmotic BBB disruption, which increases paracellular uptake of chemotherapeutics and has been applied to patients with brain tumors [22]. Paracellular permeability is regulated by cadherin-mediated adherens junctions, which can be transiently modulated by cadherin peptides such as ADTC5 (Cyclo(1,7)Ac-CDTPPVC-NH_2_), which we recently developed and validated [23]. ADTC5 binds to cadherins, transiently disrupts cadherin-cadherin interactions, and opens the BBB for 2-4 hr, thereby facilitating the delivery of molecules up to 150 kDa [24]. Given that the human ApoE protein has a molecular weight of ∼34 kDa, we hypothesized that ADTC5 could enhance the brain deposition of rhApoE2. To assess the efficacy of rhApoE2 delivery by ADTC5 in the mouse brain, rhApoE2 was labeled with IRdye800CW, and near-infrared fluorescence (NIRF) imaging was used to measure its deposition in the brain [24]. The dye that labeled rhApoE2 had a molecular weight of approximately 1 kDa, and excess dye was removed using a Pierce Zeba desalting spin column (Fig. 3A). The purity of dye-conjugated rhApoE2 was assessed by SDS-PAGE and further confirmed by Western blotting with an anti-ApoE antibody (Fig. 3B-3C). The IRdye-rhApoE2 was subsequently administered via tail vein injection to 4-6-month-old female and male hApoE4KI mice. 40-60 min post-injection, mice were euthanized using CO_2_ inhalation, and transcardial perfusion was performed with PBS+Tween-20 to eliminate IRdye-rhApoE2 and residual dye (resi-IRdye) from the brain microvessels, thereby minimizing false-positive effects. NIRF signals from all scan levels were significantly higher in mice treated with IRdye-rhApoE2 in combination with ADTC5 than in control groups (Fig. 3D-3E). The delivery of rhApoE2 was further confirmed by comparing brain tissue homogenate to a standard containing free dye using SDS-PAGE (Fig. 3F). Deposition of IRdye-rhApoE2 in mouse brains was quantified by the summed fluorescence intensity of each brain (Fig. 3G). In conclusion, rhApoE2 can be successfully delivered into the brains of hApoE4KI mice via tail vein injection facilitated by ADTC5.

**Fig. 3.**
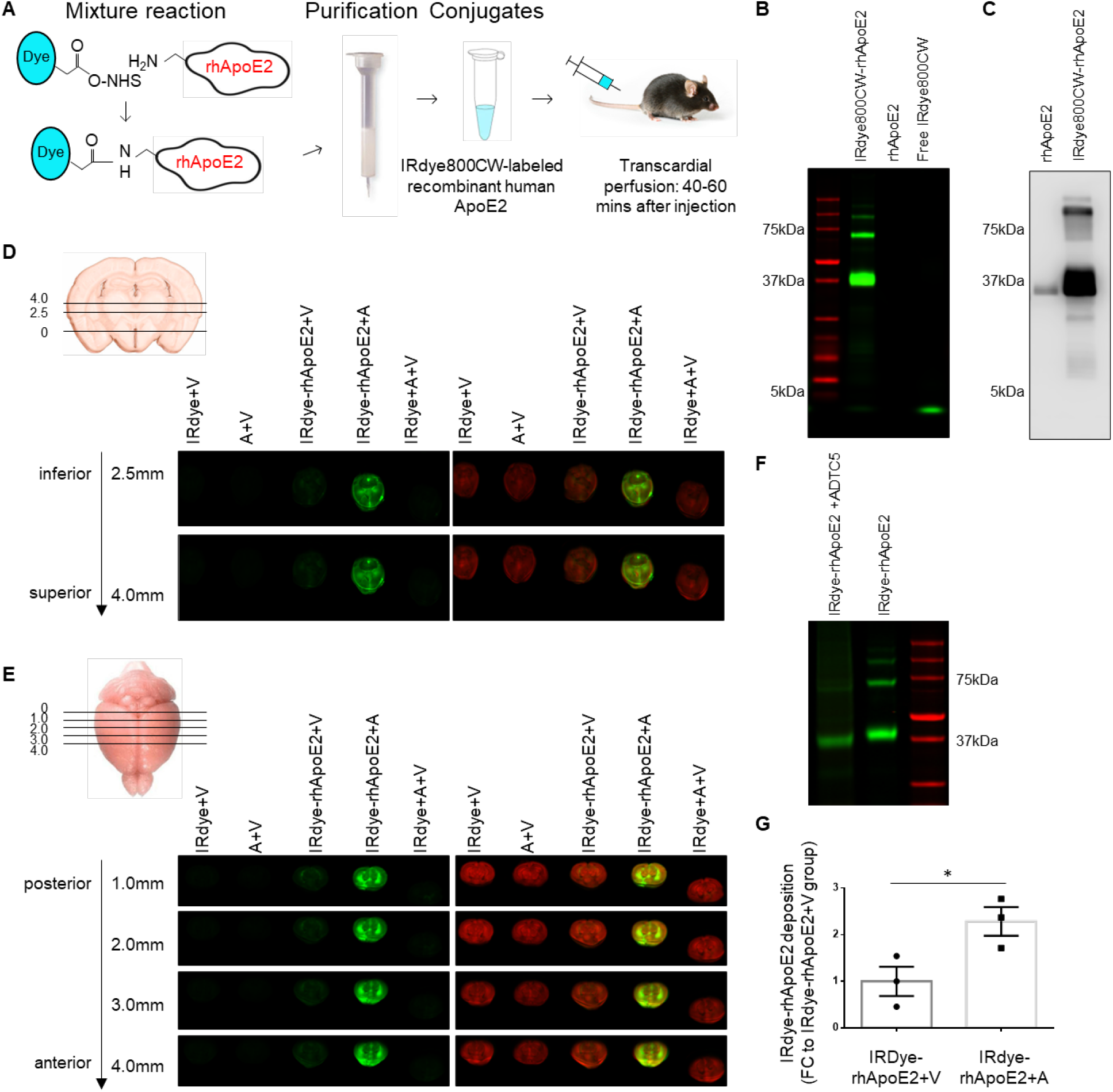
ADTC5 facilitated brain delivery of rhApoE2 in adult hApoE4KI mice. (**A**) Illustration of the generation and purification of IRdye800CW-conjugated rhApoE2. (**B-C**) The purity of the IRdye-rhApoE2 conjugate was assessed by SDS-PAGE and near-infrared fluorescence (NIRF) scanning on an Odyssey CLx imaging system. The purity of the conjugate was further confirmed by anti-ApoE antibody in WB. (**D-E**) IRdye-rhApoE2, with or without ADTC5, was injected via the tail vein into 4-6-month-old hApoE4KI mice, followed by transcardial perfusion 40-60 min post-injection to remove IRdye-rhApoE2 from brain microvessels. Mouse brains were immediately extracted and scanned on the Odyssey CLx imaging system. NIRF imaging data showed that co-administration of IRdye-rhApoE2 with ADTC5 led to significant deposition of IRdye-rhApoE2 throughout the brain; by contrast, when administered without ADTC5, IRdye-rhApoE2 deposition in the brain was nearly undetectable. (**F**) The presence of the delivered IRdye-rhApoE2 in the brain tissue homogenate was further confirmed by SDS-PAGE compared to pure IRdye-rhApoE2. (**G**) Deposition of IRdye-rhApoE2 in mouse brains was quantified by the summed fluorescence intensity of each brain. \**p*<0.05. Data are presented as the group mean ± SEM. Group comparisons were analyzed using Student’s t-test. n= 3 mice/group.

### Four weeks of weekly intravenous doese of rhApoE2 with ADTC5 increased cortical HK2 expression, hexokinase enzymatic activity, and synaptosomal vesicle release in middle-aged hApoE4KI mice

To evaluate the therapeutic impact of rhApoE2 delivery on brain changes associated with ApoE4, 15-18-month-old hApoE3KI and hApoE4KI mice of both sexes were administered pure rhApoE2 along with ADTC5, ADTC5 alone, or vehicle alone via tail vein injection once weekly for 4 weeks. Brains were harvested 24 hr after the last injection, and cortical tissues were used for subsequent experiments (Fig. 4A). Body weights of all groups before and after treatment were comparable (Fig. S3A). Western blot analysis showed no significant change in HK1 in mouse cortical lysates (data not shown), while a marked increase in HK2 was observed in mice treated with rhApoE2 along with ADTC5, compared with those treated with ADTC5 alone (Fig. 4B). Hexokinase activity assay also showed a significant increase in the rhApoE2+ADTC5 group compared with the ADTC5 alone group (Fig. 4C). These findings align with *in vitro* results, demonstrating the positive role of rhApoE2 administration in reversing ApoE4-associated changes *in vivo*.

**Fig. 4.**
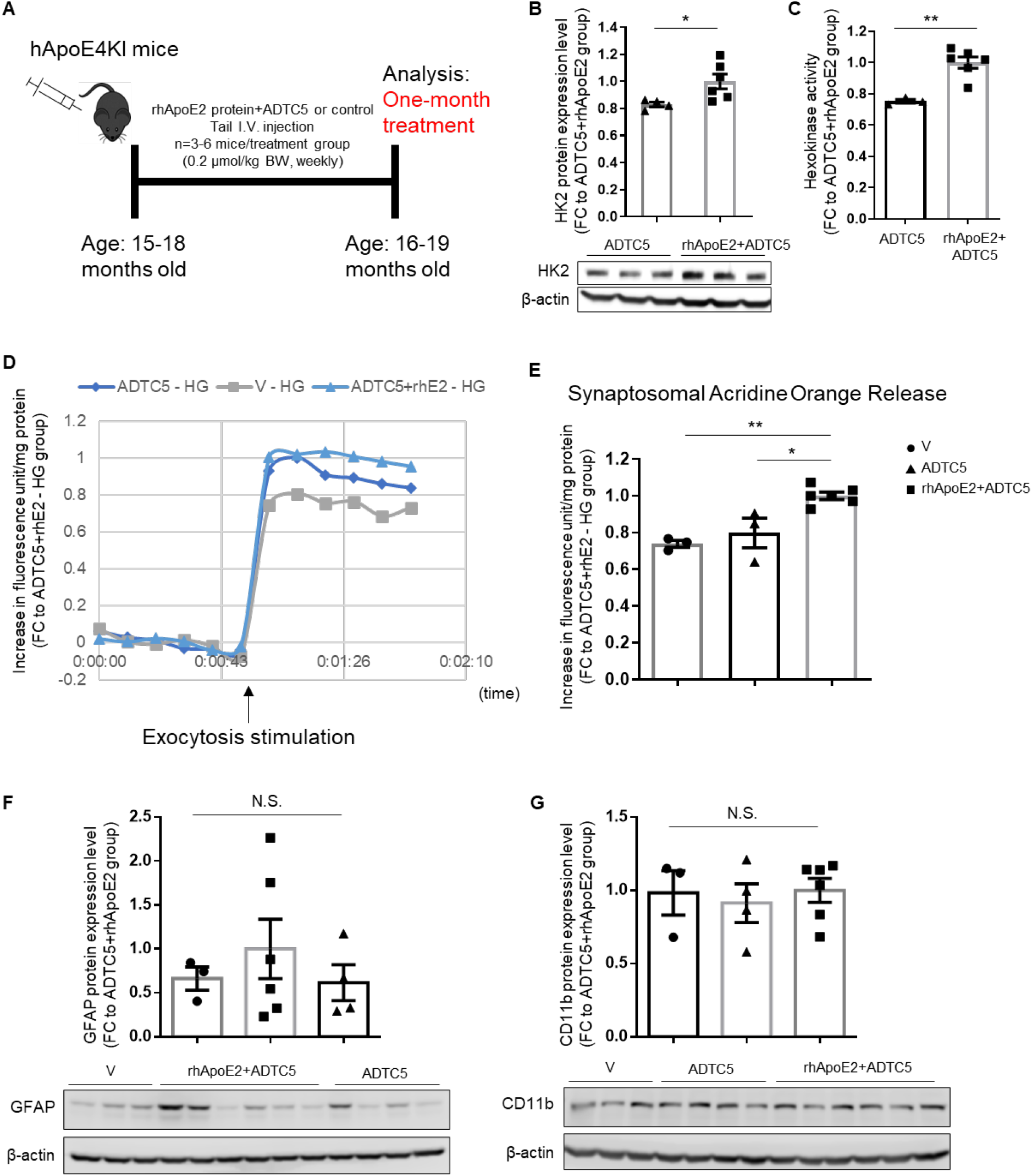
One-month intravenous doses of rhApoE2 with ADTC5 increased cortical HK2 expression, HK enzymatic activity, and synaptosomal vesicle release activity in middle-aged hApoE4KI mice. (**A**) Schematic experimental design for assessing neural responses to one-month intravenous administration of rhApoE2 in 15-18-month-old hApoE4KI mice. (**B-C**) Western blot data showed that rhApoE2 increased HK2 protein expression and HK enzymatic activity in the cortex lysate of hApoE4KI mice. (**D-E**) Cortical synaptosomes were immediately isolated after euthanasia, and synaptic vesicle release activity was evaluated using the Acridine orange assay. Results showed that synaptosomal release activity in response to KCl-induced exocytosis was enhanced in hApoE4KI mice treated with rhApoE2 compared to the V group or the ADTC5 group under high-glucose conditions. (**F-G**) No prominent gliosis was observed, evidenced by comparable levels of GFAP and CD11b expression in all treatment groups. \**p*<0.05, \*\**p*<0.01. N.S., non-significant. Data are presented as the group mean ± SEM. Group comparisons were analyzed using one-way ANOVA with Tukey’s post hoc or Student’s t-test. n = 3-6 mice/group.

Brain function was assessed by measuring synaptosomal exocytosis using the acridine orange release assay. Acridine orange is a fluorescent dye widely used to study synaptosomal vesicular release activity [25]. As a weak base, it becomes protonated and accumulates in acidic synaptic vesicles when incubated with synaptosomes, where it aggregates and causes fluorescence quenching. Upon depolarization-induced exocytosis, the dye is released, deaggregates, and increases fluorescence in the solution, making it a reliable indicator of synaptic release. Previous studies have shown that hyperglycemia can impair synaptic vesicular exocytosis, in part by disrupting glycolysis-fueled V-ATPase assembly [25]. Here, we found that mice treated with rhApoE2 along with ADTC5 showed a significant increase in synaptosomal fluorescent dye release under high-glucose conditions, indicating improved synaptic function conferred by rhApoE2, particularly under hyperglycemic stress (Fig. 4D-4E). Importantly, because prolonged BBB opening could pose risks to brain health, gliosis was assessed by measuring the expression levels of the astrocytic marker GFAP, the microglial marker CD11b, and BAX, a marker of potential apoptosis. No significant changes were observed across GFAP and CD11b levels in all hApoE4KI and hApoE3KI groups (Fig. 4F-4G and Fig. S3B-S3C). BAX levels were also comparable between the ADTC5 and vehicle-treated groups (Fig. S3B-S3C). Collectively, these results demonstrate that rhApoE2 enhances brain defenses against stressors such as hyperglycemia, likely through regulation of HK2.

### Eight weeks of weekly intravenous doses of rhApoE2 with ADTC5 upregulated cortical HK2 expression, synaptosomal glycolytic activity, learning and memory performance, and downregulated endogenous ApoE4 in aged hApoE4KI mice in a sex-dependent manner

To further evaluate the *in vivo* effects of rhApoE2 on the brain, a two-month study was conducted in aged (20-21-month-old) hApoE3KI and hApoE4KI mice of both sexes. Mice were treated with intravenous administration of pure rhApoE2 along with ADTC5, ADTC5 alone, or vehicle alone via tail vein injection, once weekly for 8 weeks (Fig. 5A). Two days before the last injection, mice were subjected to behavioral evaluation of learning and memory function in novel object recognition (NOR) (Fig. S4A) and Y-maze two-trial tests (Fig. S4C). The results showed that, compared with the hApoE4KI male control group, hApoE4KI male mice treated with rhApoE2 exhibited significantly improved performance, as evidenced by increased interactions with the novel object (Fig. 5B). In contrast, there was no improvement in NOR in female mice treated with rhApoE2 compared with the female control group (Fig. 5B). The plots in Fig. 5C show a representative heat map of the animal’s head position for the group throughout the entire NOR test, with the upper item designated as the novel object. Additionally, the total distance traveled in the field was comparable among all groups, indicating that the observed difference was independent of mobility (Fig. S4B). The Y-maze two-trial test assesses a higher level of cognitive complexity compared to the one-trial spontaneous alternation test, due to the introduction of a memory retrieval period between learning and recognition (Fig. S4C) [26]. Results showed a significant increase in the number of entries and the amount of time spent in the novel arm by hApoE4KI male mice treated with rhApoE2 compared to the male control group; however, no differences were observed in the female groups (Fig. 5D). The plots in Fig. 5E show a representative heat map of the animal’s head position for the group during the first 2 min of exploration in the Y-maze, with the lower arm set as the novel arm. The total distance traveled in the maze did not differ significantly among groups, suggesting that differences in exploration were not due to mobility (Fig. S4D). In conclusion, rhApoE2 treatment improved learning and memory in hApoE4KI male mice but not in female mice.

**Fig. 5.**
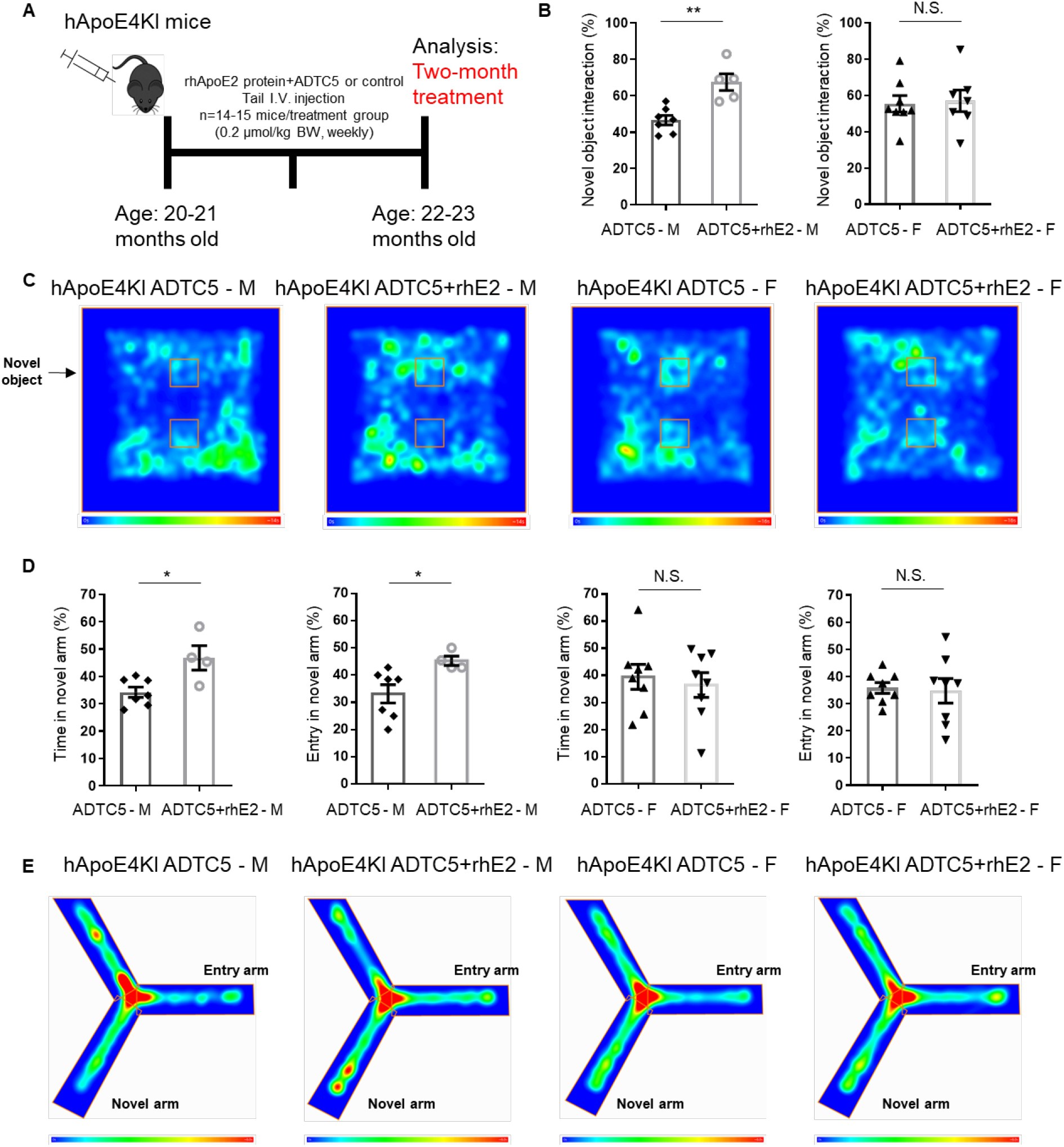
Two-month intravenous doses of rhApoE2 with ADTC5 improved learning and memory behavioral performance in aged hApoE4KI male mice. (**A**) Schematic experimental design for assessing neural responses to two-month intravenous administration of rhApoE2 in 20-21-month-old hApoE4KI mice. (**B-C**) In the novel object recognition (NOR) test, hApoE4KI male mice treated with rhApoE2 exhibited significantly greater interactions with the novel object compared to the male control group. However, rhApoE2 did not affect performance in the female groups. (**D-E**) In the Y-maze two-trial test, hApoE4KI male mice treated with rhApoE2 showed significantly greater entries into the novel arm and more total time spent in the novel arm than the male control group. Consistent with the NOR test, rhApoE2 did not affect performance in the female groups. \**p*<0.05, \*\**p*<0.01. N.S., non-significant. Data are presented as the group mean ± SEM. Group comparisons were analyzed using one-way ANOVA with Tukey’s post hoc or Student’s t-test. n = 4-8 mice/group.

To investigate potential mechanisms underlying this sex-specific effect, mouse brains were harvested 1-3 days after the last injection, and cortical tissues were analyzed in subsequent experiments. The body weights of all groups of mice remained comparable before and after treatment (Fig. S5A). Western blot results showed a significant reduction in endogenous ApoE4 in male mice treated with rhApoE2; however, no significant change was observed in female mice, although a trend toward a decrease was noted (Fig. 6A). In contrast, rhApoE2-treated female mice exhibited a significant increase in HK2 expression (Fig. 6B), whereas this effect was not observed in male mice despite a trend toward an increase (*p* = 0.1). Glycolytic function in synaptosomes was further assessed using the Seahorse XF Analyzer with the Seahorse glycolytic stress test. Consistent with the cognitive behavioral data, hApoE4KI male mice treated with rhApoE2 showed a significant increase in the basal glycolytic rate and maximum glycolytic capacity compared with the male control group. However, these outcomes were unchanged in female mice treated with rhApoE2 compared with the female control group (Fig. 6C-6D). As expected, rhApoE2 treatment upregulated PI3K/Akt signaling activity, but only in hApoE4KI male mice (Fig. 6E). Moreover, rhApoE2 markedly reduced IL-1β levels in spleen tissue lysates, again only in males (female data not shown), suggesting that male mice may have responded better to rhApoE2 treatment than female mice in reducing systemic inflammation (Fig. 6F). Similarly, no prominent gliosis was observed, as evidenced by Western blotting for GFAP, CD11b, and BAX (Fig. S5C-S5J). In summary, these results demonstrate that a two-month treatment with weekly intravenous administration of rhApoE2 was safe and effective in eliciting neural responses in aged hApoE4KI mice, potentially contributing to the improved cognitive performance observed in these mice. Interestingly, these effects were more prominent in male than in female mice, suggesting that sex may modulate brain responses to rhApoE2.

**Fig. 6.**
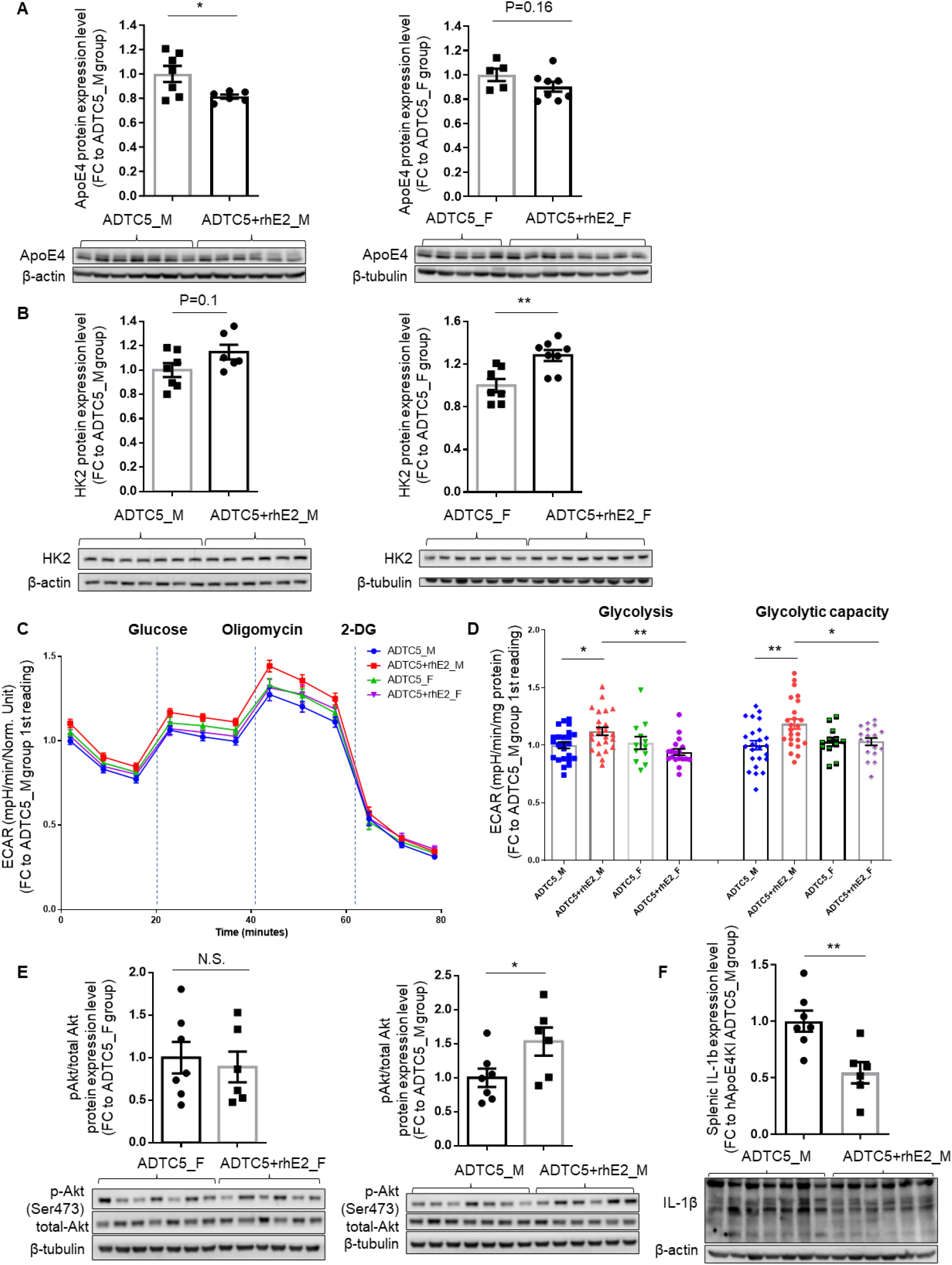
Two-month intravenous doses of rhApoE2 with ADTC5 upregulated cortical HK2 expression and synaptosomal glycolytic activity and downregulated endogenous ApoE4 in aged ApoE4KI mice in a sex-dependent manner. (**A**) Two-month intravenous exposure to rhApoE2 significantly reduced cortical expression of endogenous ApoE4 in 20-21-month-old hApoE4KI male mice but not in female mice. n = 5-8 mice/group. (**B**) Intravenous exposure to rhApoE2 significantly increased cortical HK2 expression in aged hApoE4KI female mice but not in male mice. n = 6-8 mice/group. (**C-D**) Cortical synaptosomes were isolated immediately after euthanasia, and synaptosomal glycolytic activity was measured using a Seahorse glycolytic stress test. Male mice treated with rhApoE2 had the highest levels of both basal glycolysis and maximum glycolytic capacity across groups. n = 2-4 mice/group, 6 readings per mouse. (**E**) Cortical Akt phosphorylation was significantly increased in male mice treated with rhApoE2 but not in female mice. n = 6-7 mice/group. (**F**) Similarly, male mice treated with rhApoE2 exhibited a significantly reduced IL-1β expression in spleen tissue lysate, whereas the female groups were not affected (data not shown). n = 6-7 mice/group. \**p*<0.05, \*\**p*<0.01, \*\*\**p*<0.001. Data are presented as the group mean ± SEM. Group comparisons were analyzed using one-way ANOVA with Tukey’s post hoc or Student’s t-test.

### Eight weeks of weekly intravenous doses of rhApoE2 with ADTC5 promoted DHA redistribution across phospholipids and modulated sphingolipid and galactolipid metabolism in the frontal cortex of aged hApoE4KI mice in a sex-dependent manner

Given ApoE’s critical role in lipid metabolism, particularly in light of recent studies linking ApoE4 to dyslipidemia in AD brains [27], we investigated the effects of rhApoE2 after two months of weekly intravenous administration on brain lipidomic profiles in aged hApoE4KI mice. Mouse frontal cortical tissues were collected and analyzed by untargeted lipidomics with UPLC-MS. In both male and female mice, rhApoE2 treatment altered cortical phospholipid composition, with increases and decreases in specific species (Tables S1 and S2). In male mice, significant changes were enriched in phospholipid classes, including phosphatidylethanolamine (PE), phosphatidylglycerol (PG), and phosphatidylcholine (PC) (Fig. 7A-7C), with notable redistribution of docosahexaenoic acid (DHA; 22:6), an essential omega-3 fatty acid and a building block of the brain, across phospholipid classes. Specifically, rhApoE2 induced downregulation of DHA-containing PE and concomitant upregulation of DHA-containing phosphatidylserine (DHA-PS) and DHA-PG (Fig. 7D and Table S1). Moreover, rhApoE2 significantly increased the free DHA level in male mice (Fig. 7D). In female mice, rhApoE2 treatment elicited a similar effect on DHA redistribution, but to a lesser extent, and the trend appeared to be opposite to that observed in male mice (Fig. 8A-8C). Specifically, in female mice, rhApoE2 upregulated DHA-PE species and downregulated two DHA-PA species (Fig. 8D and Table S2). Also, unlike in male mice, the free form of DHA was unaffected by rhApoE2. These data suggest a prominent role for rhApoE2 in altering ApoE4-associated lipidomic profiles by promoting metabolic dynamics and remodeling of DHA and phospholipid.

**Fig. 7.**
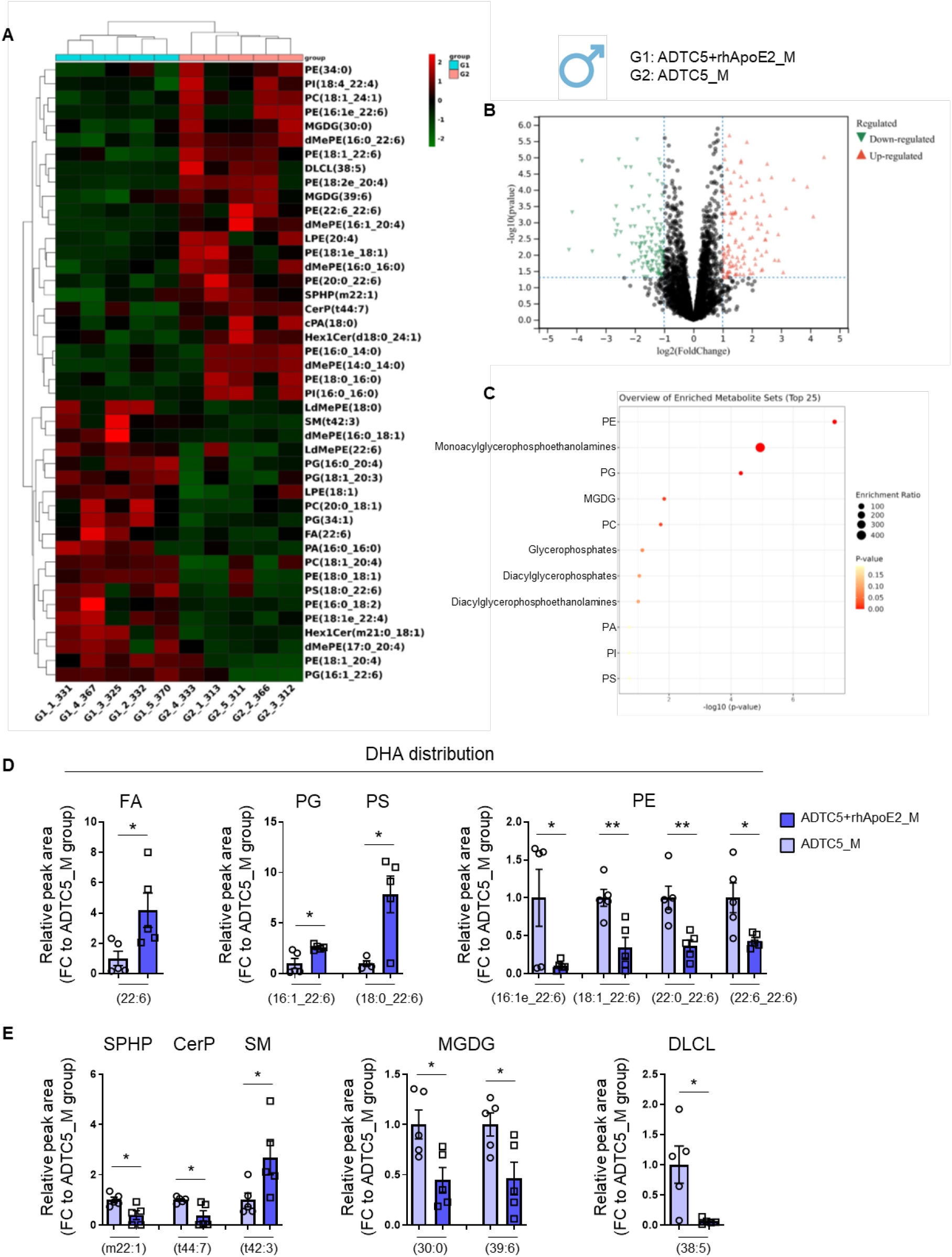
Two-month intravenous doses of rhApoE2 with ADTC5 promoted cortical DHA redistribution across phospholipids and modulated sphingolipid and galactolipid metabolism in aged hApoE4KI male mice. (**A-C**). Two-month intravenous exposure to rhApoE2 led to significant changes in the levels of phospholipids, sphingolipids, galactolipids, and cardiolipin derivatives in the frontal cortex of 20-21-month-old hApoE4KI male mice. (A) Hierarchical clustering of significantly altered lipids. (B) Volcano plot comparing Group 1 (rhApoE2+ADTC5) and Group 2 (ADTC5). Y>1.30 and X>1 were considered significant increases. Y>1.30 and X<-1 were considered significant decreases. Red indicates upregulation by rhApoE2; green indicates downregulation by rhApoE2. n = 5 mice/group. (C) Pathway enrichment analysis revealed major lipid classes that were altered by rhApoE2. (**D)** Relative abundance of DHA-containing phospholipid species in rhApoE2-treated mouse cortex compared to vehicle-treated groups. (**E**) Relative abundance of SPHP, CerP, SM, MGDG, and DLCL in rhApoE2-treated mouse cortex compared to vehicle-treated groups. n = 5 mice/group. \**p*<0.05, \*\**p*<0.01. Data are presented as the group mean ± SEM. Group comparisons were analyzed using Student’s t-test.

**Fig. 8.**
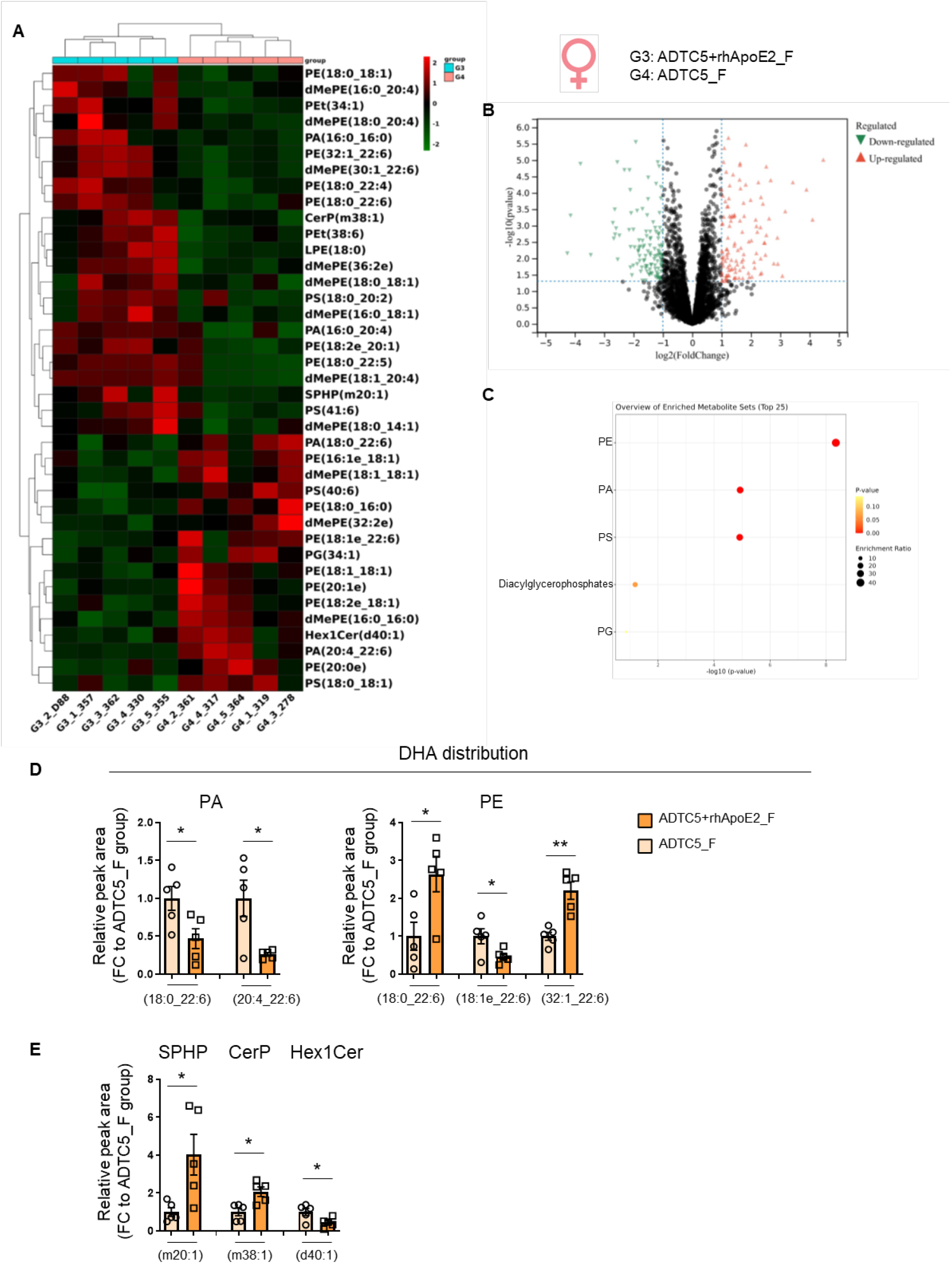
Two-month intravenous doses of rhApoE2 with ADTC5 induced a distinct pattern of cortical DHA redistribution and sphingolipid modulation in aged hApoE4KI female mice compared to male mice. (**A-C**). Two-month intravenous exposure to rhApoE2 led to significant changes in the levels of phospholipids and sphingolipids in the frontal cortex of 20-21-month-old hApoE4KI female mice. (A) Hierarchical clustering of significantly altered lipids. (B) Volcano plot comparing Group 1 (rhApoE2+ADTC5) and Group 2 (ADTC5). Y>1.30 and X>1 were considered significant increases. Y>1.30 and X<-1 were considered significant decreases. Red indicates upregulation by rhApoE2; green indicates downregulation by rhApoE2. n = 5 mice/group. (C) Pathway enrichment analysis revealed major lipid classes that were altered by rhApoE2. (**D)** Relative abundance of DHA-containing phospholipid species in rhApoE2-treated mouse cortex compared to vehicle-treated groups. (**E**) Relative abundance of SPHP, CerP, and Hex1Cer in rhApoE2-treated mouse cortex compared to vehicle-treated groups. n = 5 mice/group. \**p*<0.05, \*\**p*<0.01. Data are presented as the group mean ± SEM. Group comparisons were analyzed using Student’s t-test.

Sphingolipids were another lipid class altered by rhApoE2 in a sex-dependent manner. Major subclasses include sphingosine phosphate (SPHP) and ceramide phosphate (CerP), both of which have been linked to AD [28]. Male mice treated with rhApoE2 showed decreased levels of SPHP(m22:1) and CerP(t44:7) (Fig. 7E), whereas female mice treated with rhApoE2 showed increased levels of SPHP(m20:1) and CerP(m38:1) (Fig. 8E). In addition, male mice showed increased levels of the sphingomyelin species SM(t42:3) (Fig. 7E), though female mice showed decreased levels of the hexosylceramide species Hex1Cer(d40:1) in response to rhApoE2 (Fig. 8E). Sex differences were also observed in changes in monogalactosyldiacylglycerol (MGDG), a galactolipid class also linked to AD (Fig. 7E) [29]. Two MGDG species, MGDG(30:0) and MGDG(39:6), were significantly downregulated by rhApoE2 in male mice, whereas no changes were observed in female mice (Fig. 7E). Similarly, a dilysocardiolipin molecule, DLCL(38:5), was downregulated in male mice but not in female mice (Fig. 7E). These sex-specific changes in cortical lipid composition in response to rhApoE2 likely contributed to the observed sex differences in other brain changes elicited by rhApoE2. Collectively, these findings demonstrate that a two-month treatment with rhApoE2 via weekly intravenous administration was effective in altering brain lipidomic profiles, including phospholipids, sphingolipids, and galactolipids, in aged ApoE4 brains, with sex acting as a modulator.

## DISCUSSION

This study presents the first investigation of rhApoE2 as a potential protein therapy in human ApoE4-expressing mouse models. First and foremost, we developed and validated an expression platform to produce rhApoE2 at high yield and purity and with extensive glycosylation and sialylation that resemble human brain ApoE2 [17]. In ApoE4-expressing primary neurons, exposure to rhApoE2 elicited neuroprotective responses under both normal and neurodegenerative conditions, including upregulation of HK2 and reduction of endogenous ApoE4. To investigate whether the *in vitro* effects of rhApoE2 could be replicated *in vivo*, we first demonstrated the safe and efficient delivery of intravenously administered rhApoE2 with ADTC5 to the brains of adult hApoE4KI mice. ADTC5, a cadherin-derived peptide, transiently and reversibly increases the permeability of the paracellular pathway of the BBB by modulating E-cadherin interactions at the adherens junctions of endothelial cells, enabling the brain entry of therapeutic molecules [23]. In line with *in vitro* findings, *in vivo* studies showed that intravenous administration of rhApoE2 together with ADTC5 enhanced HK2 expression and activity, synaptic glycolysis and exocytosis, cortical lipid metabolic dynamics and DHA redistribution, and learning and memory. Intriguingly, a sex-specific effect became increasingly apparent with age, especially in aged hApoE4KI mice (20-21-month-old), with a broader and more significant positive impact in male mice. These results position rhApoE2-based therapy as a potential intervention for AD, particularly in at-risk individuals carrying ApoE4. Moreover, they provide further support to the understanding of the prominent interactions among the triad of risk factors for sporadic AD: age, sex, and ApoE [30, 31].

Our observation that rhApoE2 suppresses endogenous ApoE4 levels is particularly notable given the potential of emerging therapeutic strategies aimed at downregulating ApoE4 to prevent AD [32–34]. Attempted strategies include Cre recombinase-mediated editing, knockout of the ApoE transcription factor C/EBPβ, and ApoE-targeted immunotherapies [34–36]. Anti-ApoE antibodies were shown to decrease Aβ plaque formation by disrupting ApoE-Aβ interactions and co-deposition in APPswe/PS1ΔE9 mice [37]. Antisense oligonucleotides (ASOs) targeting human ApoE4 reduced endogenous ApoE4 by ∼50% in a mouse model of tauopathy, with attenuated neuroinflammation, sustained synaptic density, and decreased levels of the blood biomarker neurofilament light chain [32]. In addition to the ApoE4-targeted approach, ApoE2 gene therapy has also been pursued. Delivery of the ApoE2 gene via adeno-associated viral vectors by direct intracerebroventricular injection was shown to produce a notable neuroprotective effect by reducing Aβ plaque accumulation [38]. However, genome- and gene-based therapies are often limited by potential off-target effects of CRISPR/Cas9 and ASOs, overly strong immune responses, and a high risk of chromosomal disruption and insertional mutagenesis [39–41].

To our knowledge, the present study is the first to investigate the therapeutic potential of a human ApoE2 protein-based therapy to reduce AD risk or intervene in AD progression. Our group recently reported that the three ApoE isoforms in human brains undergo varying degrees of sialylation, the posttranslational process in which sialic acid moieties are attached to the terminus of glycan chains on ApoE, with ApoE2 most abundantly sialylated and ApoE4 least sialylated. In contrast, ApoE in the mouse brain was found to be minimally sialylated. These human versus rodent differences suggest that sialic acid moieties on human ApoE may play a significant role in modulating the physiological function of ApoE in the human brain [17]. For example, ApoE sialylation has been shown to influence ApoE interactions with Aβ peptides; enzymatic removal of sialic acids from ApoE2 enhanced its binding affinity for Aβ peptides and promoted Aβ fibrillation [17]. Sialic acids are most concentrated in the brain, where they critically support neuronal sprouting, synaptic plasticity, axon myelination, myelin stability, and remodeling of neuronal connections [42]. They also serve as endogenous ligands for microglial Siglecs (sialic acid-binding immunoglobulin-type lectins), such as CD33, helping maintain microglial homeostasis, while dysfunction in this signaling pathway is linked to neuroinflammation in AD [43]. These findings led us to believe that the most extensive sialylation in ApoE2 confers a structural advantage that helps protect the brain against neurodegenerative risks, thereby underpinning ApoE2-mediated resistance to AD. Thus, to properly evaluate the therapeutic potential of an ApoE2 protein-focused approach, the most critical task we undertook was to develop a method to produce ApoE2 protein with a sialylation profile mirroring that of endogenous ApoE2 in human brains. Regrettably, prokaryotic system-derived ApoE, e.g., from *E. coli*, commonly used in prior studies, lacks these essential PTMs, raising concerns about the human relevance of the findings reported in those studies [16].

AD has long been recognized as a metabolic disease, with glucose hypometabolism established as a prominent anomaly that emerges during the prodromal stage of AD [2]. Reduced glycolytic flux was found to correlate closely with the extent of plaque and tangle deposition in specific brain regions of AD patients [1]. Diminished glycolytic activity in AD was further demonstrated by significantly lower levels of key glycolytic intermediates in the CSF of AD patients [44]. Moreover, brain hyperglycemia and impaired glucose metabolism also provide a mechanistic link between AD and diabetes [25, 45]. Collectively, glycolytic improvement by rhApoE2 has significant mechanistic implications for enhancing brain resilience against AD. Our group has recently reported that ApoE isoforms differentially modulate brain bioenergetics with ApoE2 upregulating and ApoE4 downregulating neuronal HK expression and glycolytic activity compared to ApoE3 [14, 46, 47]. These ApoE-specific glycolytic status appears to strongly influence neuronal health and aging [15]. Here, we demonstrate that introducing rhApoE2 sialoprotein increased HK expression and glycolytic activity in both in *vitro* and *in vivo* ApoE4-expressing models. Our observations align with the existing literature, which indicates that neurons with higher glycolytic activity are more resistant to neurotoxicity, suggesting HK-driven glycolysis as a promising therapeutic mechanism to enhance brain resilience against AD [48].

Lipid dyshomeostasis has also been widely implicated in the pathophysiology of AD in both the brain and the peripheral circulation. A number of studies have reported an inverse association between polyunsaturated fatty acid (PUFA) supplementation, particularly omega-3 fatty acids, and AD risk [49–51]. In the present study, two-month treatment with rhApoE2 improved the cortical lipidomic profile in aged ApoE4 mice in a sex-dependent manner. In the male cortex, rhApoE2 increased levels of free DHA, PG-DHA, and PS-DHA, which could have derived from PE-DHA, as several PE-DHA species were significantly downregulated by rhApoE2. DHA constitutes over 90% of omega-3 PUFAs in the brain and plays an indispensable role in brain structure and function. DHA exists in the brain in two primary forms: free DHA and esterified DHA incorporated into membrane phospholipids. The free unesterified DHA represents a smaller, more transient pool, often released from esterified forms in phospholipids by phospholipases, and it provides anti-inflammatory and neuroprotective effects. Esterified DHA is the predominant form, and its distribution in phospholipids is dynamically regulated by the phospholipid remodeling process or the Lands’ cycle, which enables rapid DHA redistribution to support membrane fluidity and neuronal function. In the brain, PE serves as the primary structural reservoir for DHA. The redistribution of DHA from PE to PG and PS may reflect a metabolic shift that facilitates activation of signaling pathways that promote mitochondrial function and neuronal survival. For instance, increased DHA in the PS pool has been shown to inhibit neuronal death under challenged conditions by enhancing the activity of key survival kinase pathways such as PI3K/Akt, Raf-1, and PKC [52]. Interestingly, DHA recycling among phospholipids was also observed in female ApoE4 brains, but in a direction somewhat opposite to that in male brains. Specifically, the two-month rhApoE2 treatment significantly increased PE-DHA levels while reducing PA-DHA levels. These changes may indicate that the brain is shifting its DHA metabolism from the PA pathway to the PE pathway. Increased PE-DHA may suggest a refilling of the brain’s long-term DHA stores. Overall, it can be concluded that a relatively long-term introduction of rhApoE2 can induce metabolic remodeling of DHA in both female and male ApoE4 brains. In the female brain, this metabolic shift appears to be more structural, aimed at restoring homeostasis. In contrast, the shift appears to be more proactive in the male brain, promoting neuroprotective signaling in response to a challenging environment.

In addition to DHA remodeling, rhApoE2 altered sphingolipid profiles in the brains of aged ApoE4 mice, with sex also having a significant influence. Sphingosine-1-phosphate (SPHP or S1P) and ceramide-1-phosphate (CerP or C1P) were both altered in female and male ApoE4 brains, despite their distinct structures. Specifically, decreased SPHP(m22:1) and CerP(t44:7), with increased sphingomyelin SM(T42:3), were observed in male ApoE4 brains following exposure to rhApoE2. S1P plays dual roles in neuroinflammation, depending in part on disease context. Elevated S1P levels contribute to neuroinflammation across multiple conditions [53]. Indeed, several S1P receptor functional antagonists, including Fingolimod, are FDA-approved immunomodulating drugs used to treat relapsing forms of multiple sclerosis [54]. They work in part by inhibiting lymphocyte infiltration into the brain and reducing glial reactivity, thereby dampening neuroinflammation. These modulators have also been noted for their potential to improve cognitive conditions associated with AD [55]. Similar to S1P, C1P is also a bioactive sphingolipid that mediates neuroimmune responses and neuronal survival. A recent study found that total C1P levels were significantly higher in AD brains with high plaque density than in control brains and CERAD-B (classified as an intermediate probability of AD, Braak stage 1–2) [56] brains with moderate plaque density, suggesting a contribution to AD pathology [28]. Sphingomyelin (SM) is the primary structural sphingolipid of the myelin sheath and is thus crucial for neuronal communication and cognitive processes. Disruptions in SM metabolism have been implicated in various neurological conditions, including SM accumulation in Niemann-Pick disease type A/B and decreased SM in AD and ApoE4 brains [57, 58]. Treatments aimed at reversing these changes, such as those that block sphingomyelinase (SMase), the enzyme that degrades SM, are currently under investigation for their efficacy in slowing neurodegeneration [59]. In light of these earlier reports, decreases in S1P and C1P, along with an increase in SM, may indicate that rhApoE2 promoted a shift in sphingolipid metabolism in aged male ApoE4 brains, thereby restoring SM homeostasis by reducing SM breakdown into S1P and C1P. Decreased S1P and C1P may also reflect a healthier brain state with lower inflammatory activity. In contrast to male brains, female brains showed increases in two distinct species, SPHP(m20:1) and CerP(m38:1), along with a decrease in Hex1Cer(d40:1), in response to rhApoE2. Hex1Cer, or hexosylceramide, is a type of complex glycosphingolipid that has been found to accumulate to toxic levels in various neurodegenerative diseases [60]. These concomitant changes may indicate that in aged female ApoE4 brains, rhApoE2 promoted the redirection of sphingolipid flux by reducing ceramide accumulation and increasing signaling molecules such as S1P and C1P, thereby promoting brain activities that could be beneficial for brain defense against neurodegenerative risks.

The third lipid class altered by rhApoE2 exclusively in male ApoE4 brains is the galactolipid class. Specifically, two species of monogalactosyldiacylglycerol (MGDG) were downregulated by rhApoE2 in the brains of male ApoE4 mice but not in female mice. In mammalian brains, MGDGs have historically been considered minor and present at much lower levels than galactosylceramides (cerebrosides); however, recent studies indicate that they may significantly influence brain health and disease. An extensive analysis of postmortem human brain samples from three cohorts of 288 participants found that many MGDG molecular species, as well as total MGDG levels, were significantly elevated in both the gray and white matter of AD brains. Moreover, the levels of these lipid species were positively associated with clinical and pathological markers of AD severity. These findings led to the speculation that abnormal MGDG accumulation may serve as a central correlate of disease progression in AD [29]. In this context, decreases in MGDGs observed in our study may indicate another beneficial effect of rhApoE2 on lipid metabolism in male ApoE4 brains.

As discussed above, our study revealed a significant influence of sex on neural outcomes elicited by rhApoE2 treatment, with greater positive effects observed in male ApoE4 brains than in female brains. The interaction between ApoE genotype and sex as a critical factor in AD has been reported extensively in the literature, as summarized in our recent review [31, 61]. First, AD has a significant sex bias, with women being the primary victims of the disease and accounting for nearly two-thirds of all AD cases [62]. As the most significant genetic risk factor for AD, the ApoE ε4 allele has a disproportionately greater impact on women, increasing not only the risk of developing AD but also the severity of neuropathology, the extent of cognitive decline, and reducing treatment efficacy [31]. For instance, it was reported that carrying a single ApoE ε4 allele in women was sufficient to shift the age-related risk curve by approximately five years [63], conferring a disease risk comparable to that of men carrying two ApoE ε4 alleles [64]. On the other hand, Cox proportional hazards analysis of the ApoE-sex interaction on conversion risk to AD highlighted a more pronounced protective role of ApoE ε2 in male than in female subjects [30], which aligns with our findings. Based on the lipidomics data presented in this study, we speculate that the observed sex differences in neural responses to rhApoE2 likely reflect sex-specific differences prior to treatment, suggesting that the mechanisms underlying the increased AD risk conferred by ApoE4 differ between female and male brains. Future research should focus on elucidating the sex-specific risk associated with ApoE4 in aging brains, which will provide important insights for advancing precision medicine and for developing sex-specific strategies for AD prevention and treatment.

In conclusion, this work represents the first attempt to investigate the translational potential of a human ApoE2 protein-based approach for the prevention and intervention of AD. The significance and novelty of the study are several-fold. First, it focused on a highly glycosylated and sialylated form of recombinant ApoE2 that resembles the endogenous ApoE2 expressed in human brains. Second, it presented a novel method for the safe and effective delivery of rhApoE2 to the brain, without the potential side effects associated with AAV-based gene delivery. Third, it provided compelling evidence for the therapeutic potential of rhApoE2 as a promising approach that could enhance the brain’s metabolic adaptability and robustness against age-related neurodegenerative risks that could lead to AD in at-risk individuals, such as ApoE4 carriers. This neuroprotection-focused strategy is expected to be advantageous because it potentially avoids the off-target risks associated with traditional pathology-based strategies, such as anti-amyloid therapies, whose clinical efficacy in the treatment of AD remains debated.

## MATERIALS AND METHODS

### Sex as a biological variable

Both male and female animals were included in this study, and sex was considered a biological variable in data analysis.

### Study design

The goal of this study was to evaluate neuronal and brain responses to recombinant human ApoE2 sialoprotein in humanized ApoE4 knock-in mouse models, focusing on HK expression, glycolytic metabolism, synaptosomal exocytosis, changes in lipid profiles, and learning and memory. Sialylated rhApoE2 was produced and characterized using the previously established method [17].

*In vitro* experiments were performed on primary cortical neurons isolated from hApoE4KI mice on postnatal days 0-2. Neuronal responses to rhApoE2 were evaluated under both normal and neurotoxic conditions by exposing neurons to culture media containing H_2_O_2_, oligomeric Aβ, or conditioned media collected from LPS-activated microglia. Neuronal viability, HK expression, Akt activation, and endogenous ApoE4 protein levels were evaluated in at least two distinct cultures.

*In vivo* experiments were conducted in adult hApoE4KI mice of both sexes. Three separate studies were performed. The first study was conducted in 4-6-month-old hApoE4KI mice to evaluate the safety and efficacy of brain delivery of rhApoE2 facilitated by the BBB-modulating peptide ADTC5. The second study was conducted in 15-18-month-old hApoE4KI mice. rhApoE2 or vehicle controls were administered intravenously via tail-vein injections weekly, with or without ADTC5, for 4 weeks. Brain responses were analyzed to evaluate the long-term safety profile of ADTC5-facilitated brain delivery, as well as the effects of rhApoE2 on HK expression and activity and on synaptosomal release activity under hyperglycemic conditions. The third study was conducted in 20-21-month-old hApoE4KI mice to evaluate the long-term effects of rhApoE2 in aged ApoE4-expressing brains. Similar to the second study, rhApoE2 or vehicle controls were administered weekly via tail vein injections, with or without ADTC5, for 8 weeks. Before tissue harvest, mice were evaluated for learning and memory using Y-maze two-trial and novel object recognition behavioral tests. Brain responses were analyzed to assess the effects of rhApoE2 on HK expression, synaptosomal glycolytic flux, Akt activation, lipidomic profiles, and inflammatory markers in peripheral tissues.

## Animals

Animal Use Statement (AUS # 220-08) was approved by the Institutional Animal Care and Use Committee at the University of Kansas. Procedures were conducted in accordance with institutional guidelines. The humanized ApoE knock-in (hApoEKI) mice were purchased from the Jackson Laboratory (stock #029018, #027894) (Bar Harbor, ME, USA). hApoEKI mice express human ApoE isoforms from the mouse endogenous ApoE locus at a physiological level [65]. Mice were bred and housed at a consistent room temperature and humidity under a standard 12 hr/12 hr light-dark cycle. Both male and female mice were included in the study. Animals were used across three major experimental applications: (1) generation of primary neuronal cultures from P0-2 neonatal mice, (2) validation of brain delivery of IRdye-conjugated rhApoE2 with or without ADTC5, and (3) evaluation of rhApoE2 therapeutic effects in middle-aged and aged hApoE4KI mice. Mice were humanely euthanized through carbon dioxide inhalation. For validation of brain delivery of rhApoE2, transcardial perfusion was performed to remove residual intravascular signal before tissue harvest. Brain tissues were either used immediately to ensure synaptosome viability or harvested, snap-frozen, and stored at -80 °C for subsequent biochemical analyses.

### Production and purification of rhApoE2

Recombinant human ApoE2 protein was produced as previously described [17]. Briefly, FreeStyle 293-F cells (Thermo Fisher Scientific) were used as the protein expression system and transfected with pcDNA3.1(-)-hApoE2-StrepII using the recommended 293fectin transfection reagent according to the manufacturer’s instructions. The suspension cells were cultured with shaking at 125 rpm at 37 °C in 8% CO2 atmosphere and maintained for 4 days before the medium was harvested and concentrated by centrifuge using a 10 kDa molecular weight cutoff (MWCO) filter. The rhApoE2 was purified using a Gravity flow Strep-TactinXT Superflow high-capacity column (IBA Lifesciences, Germany). The protein was further concentrated by centrifugation at 4 °C to approximately 8-10 μg/μL. Protein concentration was determined with the Pierce BCA assay kit (Thermo Fisher Scientific). Protein purity was assessed using SDS-PAGE and verified with InstantBlue Coomassie protein stain. Protein integrity was further assessed by Western blot, probed with an anti-ApoE antibody. The final rhApoE2 protein was then aliquoted and stored at - 80 °C to avoid freeze-thaw cycles.

### Brain delivery of rhApoE2 in adult hApoE4KI mice

The ADTC5 peptide used in this study was generously provided by Dr. Teruna Siahaan from the Department of Pharmaceutical Chemistry at the School of Pharmacy, University of Kansas. Male and female hApoE4KI mice aged 4-6 months were divided into five groups, each consisting of three mice, and injected with: (1) IRdye-rhApoE2 (0.2 μmol/kg) and ADTC5 peptide (13 μmol/kg), (2) IRdye-rhApoE2 (0.2 μmol/kg) alone, (3) ADTC5 peptide (13 μmol/kg) alone, (4) residual IRDye (3 nmol/kg) alone, and (5) residual IRDye (3 nmol/kg) and ADTC5 peptide (13 μmol/kg). IRdye-rhApoE2 was dissolved in PBS and possesses a molecular weight of ∼37 kDa. The amount of residual free dye in the IRdye-rhApoE2 solution was estimated by comparing the signal ratio to that of IRdye-rhApoE2 in SDS-PAGE, lane 1 (Fig. 3B). The mice were euthanized by CO_2_ inhalation 40-60 min after injection, and the brains were perfused transcardially with PBS containing 0.5% Tween-20. The brains were then imaged using the Licor Odyssey CLx (Licor, Lincoln, NE). The imaging displayed fluorescence throughout the entire brain. 4 optical sections were taken at 1.0 mm increments, beginning at the transverse fissure and extending to a depth of 4 mm in the coronal plane. Results were quantified using the summed fluorescence intensity of each brain. Another 2 optical sections were taken, beginning at the bottom surface of the brain and extending to a depth of 4 mm in the horizontal plane. The fluorescence intensity was calculated by summing the values of all optical sections for each brain [24].

### Four weeks of treatment with rhApoE2 in middle-aged hApoE4KI mice

For the four-week treatment, 15-18-month-old hApoE3KI and hApoE4KI mice (n = 3-6 mice/group, both sexes) were randomly assigned to receive (1) rhApoE2 (0.2 μmol/kg) and ADTC5 peptide (10 μmol/kg), (2) ADTC5 peptide (10 μmol/kg) alone, or (3) vehicle control (PBS and buffer), via tail vein injections. Treatment was conducted weekly for 4 weeks. Mice were euthanized and analyzed at 16-19 months of age. Endpoint measurements included body weight, hexokinase expression and activity, synaptosomal activity, and potentially induced neurotoxicity.

### Eight weeks of treatment with rhApoE2 in aged hApoE4KI mice

For the eight-week treatment, 20-21-month-old hApoE3KI (n = 5-10 mice/group) and hApoE4KI mice (n = 14-15 mice/group, both sexes) were assigned to the same treatment groups and dosage regimens as described above. Mice were treated for 8 weeks and analyzed at 22-23 months of age. Outcome measurements included body weight, HK expression and activity, endogenous ApoE4 expression in cortical tissue, potential neurotoxicity, splenic inflammatory markers, PI3K/Akt signaling activity in cortical tissue, synaptosomal HK activity and glycolysis, spatial recognition memory, and lipidomic analysis of serum samples.

### Synaptosome vesicle release test

Experiments were performed as described with some modifications [25]. The dye solutions were prepared fresh and kept on ice, with light blocked throughout the procedure. Samples were prepared from a synaptosome stock at a final glucose concentration of 2.5 mM or 10 mM and a final protein concentration of around 0.2 mg/mL. CaCl_2_ and dye were added to the samples just before incubation. The final concentration of CaCl_2_ and acridine orange in the solution was 2 mM and 5 μM, respectively. Three to four replicates of the 100 μL sample were loaded into a 96-well opaque plate and incubated for exactly 10 min at 37 °C. After incubation, kinetic fluorescence measurements were conducted at 37 °C using a Molecular Devices SpectraMax iD3 plate reader. The excitation/emission wavelengths were set at 490 nm. Initially, the baseline was recorded for 2 min to stabilize the fluorescence reading of the solution. KCl was then injected into the wells to a final concentration of 60 mM to induce exocytosis and release of the dye into the solution. Afterward, the plate was shaken, and fluorescence was remeasured kinetically for another 2 min. Wells containing dye without synaptosomes served as the negative control. Wells containing the 2x protein concentration (around 0.4 mg/mL) served as positive control. The increase in fluorescence reading from baseline after KCl injection was recorded and normalized to the total protein amount in the sample.

### Synaptosome glycolytic stress test

Synaptosome pellet samples were prepared following the above method and dissolved in Ionic Media, as previously reported [66]. To prepare 1 L of Ionic Media, HEPES (4766.044 mg), Na_2_HPO_4_(170.4 mg), D-glucose (1801.6 mg), MgCl_2_ (95.2 mg), NaHCO_3_ (252 mg), KCl (372.8 mg), and NaCl (8176 mg) were dissolved. The pH of the media was adjusted to 7.4, and the volume was brought to 1 L with double-distilled water. Subsequently, 40 ul of the resuspended synaptosomes were plated onto a Seahorse XF96 poly-D-lysine-coated plate and centrifuged at 3,220 x g for 40 min at 4°C to promote adherence of the synaptosomes to the well bottom (Agilent Technologies, Santa Clara, CA, USA). The synaptosomes were then incubated at 37 °C without CO2 for 1 hr, after which 180 μL of warm glycolytic stress test assay medium was added, prepared as XF base medium (Agilent Technologies, Santa Clara, CA, USA) containing 2 mM glutamine, pH 7.4. The samples were then immediately measured in the Seahorse XF Analyzer following successive injections of 10 mM glucose, 2 μM oligomycin, and 50 mM 2-Deoxy-D-glucose to each well. The cell culture plates were saved after the assay and used to normalize the results by measuring protein content in each well using BCA.

### Y-maze two-trial cognition test

The Y-maze two-trial test was conducted according to previous studies with modifications [26, 67]. The Y-maze apparatus was purchased from Stoelting Co., with standard arm lengths of 35 cm, arm heights of 15.5 cm, and lane widths of 5 cm (Stoelting, Wood Dale, IL). Visual cues were applied and kept consistent throughout all tests. The testing room temperature was maintained at around 22 °C, with dimmed lighting at 15-20 LUX and minimal sound disturbance. Prior to the test, mice were transferred to the behavior test room and allowed to acclimate for 1 hr. The two-trial Y-maze test consisted of 2 sessions separated by a 30 min interval. In the first trial session, one arm was blocked, and the mice were placed at the end of that arm, facing away from the maze center, allowed to explore the two other open arms for 5 min, and then returned to their cages for 30 min. In the second trial session, the block was removed, and all three arms remained open. The mice were placed at the end of one arm facing away from the maze center and allowed 5 min of free exploration. The number of entries and the amount of time spent in each arm were recorded. To prevent odor cues between trials and different mice, the maze was wiped thoroughly with 10% ethanol. The percentage of time and entries in the novel arm was calculated as the time and entries in the novel arm divided by the time and entries spent in all three arms during the first 2 min of the second testing trial. Mice with fewer than three arm entries (excluding the start arm entry) in the first minute were set as the outlier criteria and removed from the analysis. The ANY-MAZE software was used to track, record, and analyze the test (Stoelting, Wood Dale, IL).

### Novel object recognition (NOR) test

The NOR test was conducted with modifications to the previously described method [67]. The open field used in the test was a uniformly illuminated box with dimensions of 40 x 40 x 35cm (Stoelting, Wood Dale, IL), and the NOR standard accessories were purchased from Stoelting Co. to ensure comparable features and to prevent differential preference by mice (Cat. #62007s, Stoelting, Wood Dale, IL). Light was dimmed to 15-20 LUX and sound disturbance remained as low as possible. The test consisted of four sessions: habituation, familiarization, interval, and testing session. One day prior to the testing day, the mice were allowed to explore the empty open field for 10 min to acclimatize to the environment. On the day of testing, the mice were first brought to the testing room 1 hr before the test to further habituate. In the training session, the open field was introduced with two identical objects. The mice were placed in the open field facing the wall, away from the objects, and allowed to explore for 10 min with free access to the objects before being returned to their cages. After a 45 min interval, one of the objects was replaced with a novel, unfamiliar object, and the mouse’s interactions with both objects were recorded. Novel object interaction was defined as an event where a mouse’s head was within 2 cm of the object [67]. The open field was cleaned with 10% ethanol and/or Peroxigard to remove any olfactory cues. Novel object interaction (%) was calculated as the percentage of the time spent interacting with the novel object divided by the total interaction time with both objects during the entire test session [67]. The ANY-MAZE software was utilized for tracking, recording, and data analysis (Stoelting, Wood Dale, IL). Mice with less than 5 sec of total object interaction in the testing trial were excluded from the analysis.

### LC-MS untargeted lipidomics analysis

The mouse right frontal cortex was harvested, snap-frozen, and stored at -80 °C until it was sent to Creative Proteomics for lipidomics profiling analysis (CPLC10102203-1). Briefly, samples were thawed on ice, about 50 mg of sample was weighed into a tube, and 1.5 mL Chloroform: MeOH (2:1, v/v) and 0.5 mL ultrapure water were added to the sample, ground for 180 sec at 65 Hz, and vortexed for 1 min, followed by sonication for 30 min, 4°C. Then centrifuge 10 min at 3,000 rpm, 4°C, transfer the lower phase to a new tube, and dry under nitrogen. Resuspend the dried extract in 400 μL of isopropyl alcohol: MeOH (1:1, v/v); add 5 μL LPC (12:0) as an internal standard. Finally, centrifuge 10 min at 12,000 rpm, 4 °C; transfer the supernatant for LC-MS analysis. Separation is performed by UPLC (Waters). The LC system is composed of an ACQUITY UPLC BEH C18 (100 mm ×2.1 mm,1.7 μm) column. The mobile phase is composed of solvent A (60% CAN + 40% H2O + 10mM HCOONH4) and solvent B (10% CAN + 90% isopropyl alcohol + 10mM HCOONH4) with a gradient elution (0−1.0 min, 30% B; 1.0−10.5 min, 30%-100% B; 10.5−12.5min, 100% B; 12.5−12.51 min, 100%−30% B; 12.51−16 min, 30% B). The mobile phase flow rate is 0.3 mL/min. The column temperature is maintained at 40 °C, and the sample manager temperature is set at 4°C. Raw data are acquired and aligned using the Lipid Search software (Thermo) based on the m/z values and retention times of the ion signals. Ions from both ESI- and ESI+ are merged and imported into the SIMCA-P program (version 14.1) for multivariate analysis. A Principal Components Analysis (PCA) is first used as an unsupervised method for data visualization and outlier identification. Supervised regression modeling is then performed on the dataset using Partial Least Squares Discriminant Analysis (PLS-DA) or Orthogonal Partial Least Squares Discriminant Analysis (OPLS-DA) to identify potential biomarkers. The biomarkers are filtered and confirmed by combining the VIP values (VIP > 1.5), p-values (p < 0.05), and FC values (FC > 2).

### Statistical analyses

GraphPad Prism 6 (GraphPad Software, La Jolla, CA, USA) was used for statistical analyses. Data are presented as the group mean ± SEM. One-way analysis of variance (ANOVA) with Tukey’s post hoc test or Student’s t-test was used for group comparisons. A *p*-value less than 0.05 was considered statistically significant.

## Supplementary Materials

The PDF file includes:

## Materials and Methods

Figs. S1 to S5

Tables S1 to S2

## Funding

This work was supported in part by grants from the National Institutes of Health (R01AG061038, R01AG071682, R01AG082273) and internal funds from the University of Kansas.

## Author contributions

L.Z. and X.Z. designed the experiments. X.Z. performed the experiments. X.Z. and L.Z. analyzed the data. H-J.M. contributed to the development of the methodology for molecular cloning and rhApoE2 production. T.S. contributed to the development of BBB-modulating cadherin peptides, including ADTC5, which was used in the study. X.Z. and L.Z. wrote the original draft. L.Z. finalized the manuscript. All authors provided critical feedback and helped shape the research, analysis, and completion of the manuscript. All authors have read and approved the final version of the manuscript.

## Competing interests

The authors declare that they have no competing interests.

## Data and materials availability

All data associated with this study are presented in the main text or the Supplemental Materials.

## Notes

### Competing Interest Statement

The authors have declared no competing interest.

