## Supplemental Methods Figures Tables for "Recombinant sialylated ApoE2 suppresses ApoE4 and sex-specifically strengthens brain metabolism and cognition in ApoE4 mice"

### MATERIALS AND METHODS

#### Conjugation of rhApoE2 with IRdye-800CW NHS Ester

Conjugation of rhApoE2 with IRdye-800CW was performed according to the manufacturer's instructions and a previous study [1]. Briefly, the dye reacted with 5-10 mg/mL recombinant protein in 0.1 M sodium bicarbonate buffer (pH 8.5) for at least 2 hr at room temperature in the dark. Residual dye was eliminated by a Zeba Spin Desalting column with a 7 kDa molecular weight cutoff (Thermo Fisher Scientific). The purity of the IRdye-rhApoE2 was verified by SDS-PAGE and scanned (excitation = 778 nm; emission = 794 nm) using the Odyssey CLx NIFR system.

#### Generation of β-amyloid oligomers and fibrils

The methods utilized to generate oligomeric Aβ (oAβ) and fibrillar Aβ were adapted from previous work with some modifications [2]. Briefly, lyophilized Amyloid β-Protein (1-42) obtained from BACHEM (Cat. #4014447) was stored at -80 °C in a sealed container. Before reconstitution, the peptide was equilibrated at room temperature for 30 min. Then, 1,1,1,3,3,3-Hexafluoro-2-propanol (HFIP; Cat. #52517, Sigma) was added to resuspend the peptide, and the mixture was vortexed at high speed for 10 min to enhance solubilization at room temperature. The solution was then aliquoted and left to evaporate under vacuum in the fume hood overnight to prevent contamination. The resulting clear peptide film was stored desiccated at -20 °C for future use as HFIP-Aβ. To generate Aβ oligomers, the HFIP-Aβ was initially dissolved in pure sterile dimethyl sulfoxide (DMSO) through vigorous pipette mixing to achieve a concentration of 5 mM. The peptide was then resuspended in ice-cold phenol red-free Ham's F-12 cell culture medium (Caisson Laboratories) to achieve a concentration of 500 μM, followed by incubation at 37 °C overnight.

Fibrillar Aβ was generated by dissolving the 5mM Aβ in DMSO in 10 mM HCl and incubating at 37 °C overnight. The characterization and verification of the produced oAβ was performed using SDS-PAGE with the Invitrogen NuPAGE MES SDS Buffer Kit (for Bis-Tris Gels) (Thermo Fisher Scientific). Protein samples were diluted in LDS Sample Buffer without reducing reagent and loaded into each well of a 12% Bis-Tris gel without boiling. Separation of protein samples was performed under non-reducing conditions, followed by transfer onto a PVDF membrane and incubation with anti-Aβ primary antibody (1:1000, clone 6E10, Biolegend). After washing with 1x TBST, the membrane was scanned with the C-Digit Blot Scanner (LI-COR, Lincoln, NE, USA) after application of the enhanced chemiluminescence (ECL) reagent (Bio-Rad) to detect the bands.

**Phase contrast and fluorescence imaging**

Primary neuron morphological appearances were evaluated using phase contrast imaging at 14 and 21 days in vitro (DIV). Images were acquired using a Zeiss Primovert with an AxioCam Erc5s rev 2 microscope at 10× or 20× phase contrast magnification. Zen Blue Software was used for microscope control, image acquisition, image processing, and data analysis (Carl Zeiss AG, Jena, Germany).

For immunocytochemistry (ICC) staining. 14 and 21 DIV primary neurons were selected to probe for neuronal marker MAP2 in fixed neuron culture samples. Primary neurons were seeded on PDL-coated 22 mm coverslips (Neuvitro Corporation, WA, USA) with regular maintenance. Cells were first fixed with 2% paraformaldehyde (PFA) for 15 min at room temperature (Thermo Fisher Scientific). The cells were washed with room-temperature buffer for 5 min, followed by 2 incubations with permeabilization buffer 1 (PB1), 5 min each, and 1 incubation with permeabilization buffer 2 (PB2), 5 min each, at room temperature. Cells were washed once more with PB1 and then blocked with a PB1 solution containing 3% donkey serum for 20 min. The cells were washed with 1x PB1 for 5 min, then incubated overnight at 4 °C in blocking buffer with the primary antibody. The cells were washed with PB1 three times, 5 min each, and incubated with the desired fluorophore-conjugated secondary antibody in PB1 only (no serum) for 1.5-2 hr in the dark. The cells were washed with PB1 and 1x PBS, then stained in the dark with DAPI (10 μM) for at least 45 min. Coverslips were subsequently mounted and readied for imaging or stored at 4°C for long-term use. Zeiss Axio Observer, Type: EK 130X85 mot. Tango CZ. ApoTome 2 was used for epifluorescence imaging. Zen Blue Software was used for microscope control, image acquisition, image processing, and data analysis (Carl Zeiss AG, Jena, Germany). The details of antibodies and buffers used are listed in the table below.

| Description | Vendor | Product # | ICC Dilution |
|---|---|---|---|
| MAP2 Antibody | Protein tech | 17490-1-AP | 1:400 |
| Donkey anti-rabbit IgG - AlexaFluor 647 | Invitrogen | A32795 | 1:200 |
| Normal Donkey Serum | Abcam | ab7475 | 3% |

| Buffer recipe | |
|---|---|
| Buffer | PBS + 0.01% Sucrose |
| Permeabilization buffer 1 (PB1) | PBS + 0.01% Sucrose + 0.1% Saponin |
| Permeabilization buffer 2 (PB2) | PBS + 0.01% Sucrose + 0.1% Saponin + 0.01% Triton |

#### Hexokinase activity assay

The hexokinase activity assay was conducted as previously described [3]. Briefly, sample lysates were harvested as described above. Hexokinase activity was measured as the total glucose phosphorylating capacity of the lysate, based upon the reduction of $NAD^+$ through a coupled reaction with glucose-6-phosphate dehydrogenase. Results were measured spectrophotometrically by monitoring the increase in absorbance at 340 nm every 1 min for 30 min at 37 °C. The initial linear slope of the curve was used to determine ΔA/min for further calculation. The hexokinase assay solution was prepared with 13.3 mM $MgCl_2$, 0.112 M glucose, 0.55 mM adenosine 5′-triphosphate, 0.227 mM $NAD^+$, and 1 IU/mL glucose-6-phosphate dehydrogenase in 0.05 M Tris-HCl buffer (pH 8.0) as described previously [4]. After 6–8 min of incubation at room temperature to reach equilibrium, 15–20 µg of the extract was added to 150 µL assay solution to initiate the reaction. Results were normalized to total cellular protein content using the BCA assay.

**LDH release and metabolic activity assays**

The LDH Cytotoxicity Detection kit (Roche Molecular Biochemicals, Mannheim, Germany) was used to assess cell viability by measuring LDH released by nonviable cells into the medium, as per the manufacturer's instructions with minor modifications. In brief, an equal amount of medium and Cytotoxicity Detection Reagent was added to a single well on a 96-well opaque plate with a clear bottom, followed by 30 min of incubation at room temperature, while avoiding exposure to light. After the stop solution was added, the colorimetric absorbance was measured at 490 nm using a Spectrophotometer with a Molecular Devices plate reader, SpectraMax iD3. Medium from wells without cells served as the control. Higher absorbance readings indicate increased cellular death and toxicity.

For the metabolic activity assay using PrestoBlue, Briefly, 10 µL of PrestoBlue reagent was directly added to cells in 90 µL of culture medium in a 96-well plate cultured with cells. The plate was incubated at 37 °C for up to 1 hr, and fluorescence was read at Ex/Em 535/560, 590/615 nm using Molecular Devices plate reader SpectraMax iD3. Wells containing only cell culture media (no cells) were used as background control wells. Higher fluorescence units indicate greater metabolic activity.

**Mouse postnatal cortical neuronal culture and treatments**

Postnatal day 0-2 hApoE4KI mice were euthanized by decapitation. Brains were taken out and placed in warm dissection medium. Cortical tissues were separated and dissected under the upright microscope and dissociated with papain solution at 37 °C with orbital shaking at 120 rpm for 30 min followed by 0.1% w/v DNase incubation for 5 min at room temperature. Cortical tissues were washed with neutral medium twice before trituration with sterile polished Pasteur pipettes for 5-10 strikes. The cell suspension was then filtered through a 40 μm cell strainer into a conical centrifuge tube and centrifuged at 170 x g for 5 min. The cell pellet was resuspended in warm maintenance medium. The cells were then counted and plated on poly-D-lysine (Sigma-Aldrich, USA) coated culture plates (Corning, USA). The specific medium and solutions used are listed in the table below.

| | Components | Final concentration | Source |
|---|---|---|---|
| Dissection medium | HBSS (Ca2 + and Mg2 + free) | 98.5% | GIBCO, Cat. #14170-112 |
| | 100mM sodium pyruvate (100×) | 1x | GIBCO, Cat. #11360-070 |
| | 20% (wt/vol) glucose (in Milli-Q water, filter sterilized) | 0.1% | Sigma, Cat. #G8270-100G |
| Neutral medium | Dulbecco's Modified Eagle Medium | 88% | GIBCO, Cat. #11965-092 |
| | FBS (re-filtered, heat-inactivated) | 10% | GIBCO, Cat. # A3840101 |
| | 100mM sodium pyruvate (100×) | 1x | GIBCO, Cat. #11360-070 |
| | Penicillin-Streptomycin (100×) | 1x | GIBCO, Cat. #15140-122 |
| Maintenance medium | Neurobasal-A Medium | 97% | GIBCO, Cat. #12349-015 |
| | B-27™ Plus Supplement (50X) | 1x | GIBCO, Cat. # A3582801 |
| | GlutaMAX Supplement (100×) | 1x | GIBCO, Cat. #35050-061 |
| Papain solution | L-Cysteine hydrochloride | 5.5 mM | Sigma, Cat. #C7477-25G |
| | Lyophilized Papain | 2 mg/ml | Fishersci, Cat. #NC9912923 |
| | Dissection medium | | |
| DNase enzymatic solution | DNase (in Milli-Q water, filter sterilized) | 1% w/v | Sigma, Cat. #DN25-100MG |

Primary neurons were isolated from postnatal 0-2 days hApoE4KI mice, cultured and maintained as described above. Under normal conditions, neurons were treated with vehicle or rhApoE2 at 25 μg/ml or 100 μg/ml for 2 days. For neurotoxicity assays, neurons were pre-treated with 25 μg/ml rhApoE2 or vehicle for 2 days prior to neurotoxic insults. Oxidative stress was induced by treatment with 100 μM $H_2O_2$ for 30 minutes, followed by continued incubation with 25 μg/ml rhApoE2 or vehicle for up to 6 hr. Amyloid toxicity was induced by exposure to 5 μM oligomeric Aβ or vehicle in the presence or absence of prior 25 μg/ml rhApoE2 treatment for two days. For microglia-mediated neurotoxicity, conditional medium from 0.1 μg/ml (low dose) or 1 μg/ml (high dose) lipopolysaccharide (LPS) activated immortalized microglial cells (IMG) was collected after 16 hr and diluted 1:200 into primary neuron culture medium and applied to ApoE4-expressing neurons with or without prior 25 μg/ml rhApoE2 for 2 days. Following treatments, cells were harvested and subjected to protein analysis by Western blot, and cell viability and metabolic activity were assessed by LDH release and PrestoBlue assays according to the manufacturer's instructions.

**IMG mouse microglial culture and collection of conditioned medium**

IMG mouse microglial cell line was obtained from MilliporeSigma (Cat. #SCC134) and maintained according to the manufacturer's manual. Briefly, cells were cultured and maintained in high-glucose DMEM (Sigma-Aldrich, USA) supplemented with 10% FBS, 1% L-glutamine, and 1% penicillin-streptomycin. Cells were cultured in a humidified incubator at 37 ℃ with 5% $CO_2$ and passaged using standard procedure upon reaching appropriate confluency. Passages below 60 were used for all experiments. LPS was purchased from Sigma-Aldrich, Inc. (Cat. #L2630) and dissolved in PBS. To obtain LPS-activated microglial conditioned medium (MCM), IMG cells were grown on 6-well plates with 2 mL complete medium in each well and treated with 0.1 μg/mL LPS as a low dose treatment LPS-MCM(L), or 1 μg/mL LPS as a high dose treatment LPS-MCM(H). The LPS-activated MCM (LPS-MCM) medium containing the entire microglia secretome was collected after 16 hr and stored at -80 °C for future use.

**Generation and maintenance of N2a-hApoE stable cell lines**

Mouse neuro-2a (N2a) cells that stably express human ApoE isoforms (N2a-hApoE) were obtained as previously described [3]. Briefly, 2 μg of human pCMV-ApoE ε2, pCMV-ApoE ε3, or pCMV-ApoE ε4 plasmid were transfected using Lipofectamine 3000. After 48 h, the medium was replaced and supplemented with Geneticin (G418, Thermo Fisher Scientific, Waltham, MA, USA). Cells were maintained and subjected to serial dilutions in 96-well plates. Approximately twenty to thirty wells of each ApoE genotype that contained only one cell were numbered and considered as the monoclonal colony. Cells were maintained in Dulbecco's Modified Eagle Medium (DMEM, high glucose) supplemented with 10% fetal bovine serum (FBS, Thermo Fisher Scientific) containing 600 μg/mL G418 in a humidified incubator under an atmosphere of 5% $CO_2$ at 37 °C.

**Transfection of ApoE2 in N2a-hApoE4 cells**

Human ApoE2 cDNA clones expressed in the mammalian vector pCMV6-Entry with C-terminal MycDDK Tag were obtained as previously described [5]. An empty pCMV6-Entry vector was used as the control. Briefly, multiple doses of human ApoE2 cDNA were transfected into N2a-hApoE4 using jetPRIME transfection reagent according to the manufacturer's manual. Fresh complete medium was replaced 4 hr after transfection. Cells were then continuously incubated for 48–72 hr before proceeding to further experiments.

**Western blotting**

Mouse splenic tissues were homogenized using the Bullet Blender in Tissue Protein Extraction Reagent (T-PER) (Thermo Fisher Scientific) supplemented with protease and phosphatase inhibitors (PPI) (Thermo Fisher Scientific), in accordance with the manufacturer's instructions (Next Advance, Inc., NY, USA). Neuronal cells were rinsed with cold PBS (pH 7.4, Thermo Fisher Scientific) and lysed in NPER+PPI for 10–15 min on ice. Whole-cell lysates were centrifuged at 4 °C for 5 min at 1,500 × g, while splenic tissue homogenate was centrifuged at 13,000 rpm for 8 min at 4 °C. The supernatants were collected. Homogenates, synaptosomes, and cytosol fractions of 25 DIV hApoE4KI primary neurons were harvested using Syn-PER Synaptic Protein Extraction Reagent with PPI according to the manual (Thermo Fisher Scientific, USA). The protein concentration was determined by the BCA protein assay kit (Thermo Fisher Scientific). Samples were then diluted in Laemmli sample buffer (Bio-Rad, Hercules, CA, USA) with 2-mercaptoethanol (Bio-Rad) and boiled at 95 °C for 5 min. An equal amount of total protein was separated by 10% SDS-PAGE. For protein purification evaluation, the gel was incubated with InstantBlue Coomassie protein stain for 2 hr at room temperature. Otherwise, the separated proteins were then transferred onto 0.2 µm pore-sized PVDF membranes (Bio-Rad). The membrane was blocked with 5% non-fat milk in TBS (100 mL 10× TBS (200 mM Tris, 1.5 M NaCl), 900 mL ddH2O, pH 7.6) for one hour at room temperature, followed by incubation with primary antibody at 4 °C overnight. The membranes were then washed with TBST [100 mL 10× TBS (200 mM Tris, 1.5 M NaCl), 10 mL 10% Tween-20, 890 mL ddH2O, pH 7.6], followed by hybridization with the horseradish peroxidase (HRP)-conjugated secondary antibody. After washing with TBST again, the bands on the membrane were detected using a C-Digit Blot Scanner (LI-COR, Lincoln, NE, USA) after application of the ECL reagent (Bio-Rad). Quantification was performed using Image Studio Version 4.0 imaging software, with normalization to the internal loading control protein. The following primary antibodies were applied: goat anti-Apolipoprotein E (1:4000, EMD Millipore, Burlington, MA, USA), mouse anti-Apolipoprotein E4 (1:2000, Novus Biologicals, CO, USA), rabbit anti-hexokinase I (1:2000, Cell Signaling Technology, Danvers, MA, USA), rabbit anti-hexokinase II (1:1500, Cell Signaling Technology), rabbit anti-Phospho-Akt (Ser473) (1:1000, Cell Signaling Technology), rabbit anti-Akt (pan) (1:1000, Cell Signaling Technology), synaptophysin (1:1000, Cell Signaling Technology), NeuN (1:1000, Cell Signaling Technology), anti-GFAP (1:2500, Calbiochem), anti-CD11b (1:1000, Abcam), BAX (1:1000, Cell Signaling Technology), and HRP anti-β-actin (1:5000, BioLegend, San Diego, CA, USA. The secondary antibodies included goat anti-Mouse, HRP (1:5000, Thermo Fisher Scientific), goat anti-rabbit, HRP (1:5000, Thermo Fisher Scientific), goat anti-Rat (1:5000, Cell Signaling Technology) and rabbit anti-goat, HRP (1:3000, Thermo Fisher Scientific).

**Synaptosome preparation**

Synaptosome samples were prepared and confirmed as previously described [6]. Briefly, Syn-PER™ Synaptic Protein Extraction Reagent (Cat. #87793) was used to isolate synaptosomes according to the manufacturer's instructions with some modifications. 30-50 mg of right frontal cortical brain tissues were minced and placed in 1 mL Syn-PER reagent and 10 μL of 100x Halt Protease and Phosphatase Inhibitor cocktail. A chilled Dounce pestle was used to homogenize the tissues using 10-15 slow strokes. The homogenates were centrifuged at 1,200 x g for 10 min at

4 °C. An aliquot of the supernatant was stored at -80 °C for further biochemical experiments as the homogenate fraction. The remaining supernatant was centrifuged again for 20 min, at 15,000 x g at 4 °C. The resulting supernatant was collected as the cytosolic fraction and stored at -80 °C. The remaining pellet was resuspended in synaptosome buffer to yield the synaptosome fraction.

Supplementary Figure 1

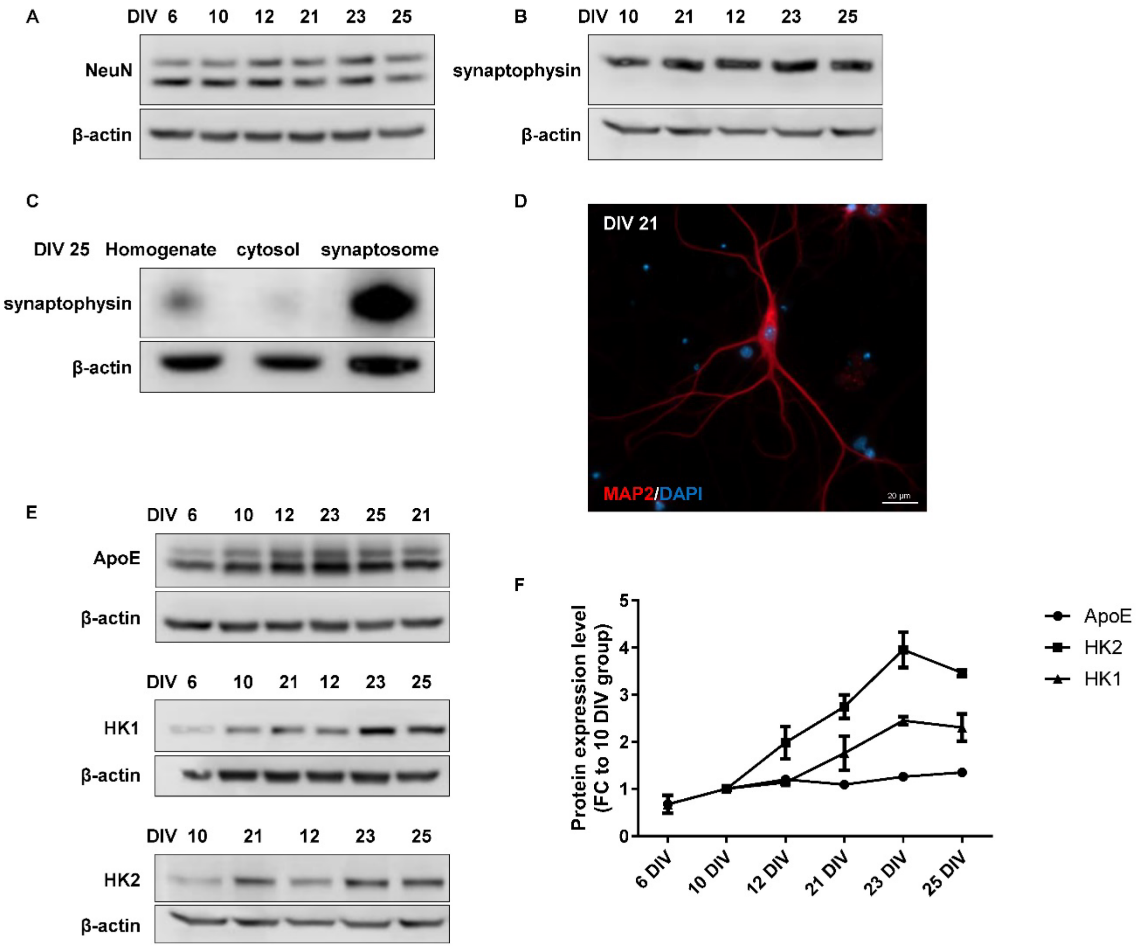

**Fig. S1. Characterization of primary cortical neuronal cultures prepared from hApoE4KI P0-2 mice.** (**A-C**) Neuronal cultures were confirmed by robust expression of the neuronal protein markers NeuN and synaptophysin. (**D**) Neuronal cultures were confirmed by MAP2 staining of neuronal cell bodies and processes. (**E-F**) ApoE, HK1, and HK2 were all expressed in neuronal cultures, with the highest expression at DIV 23.

**Supplementary Figure 2**

A

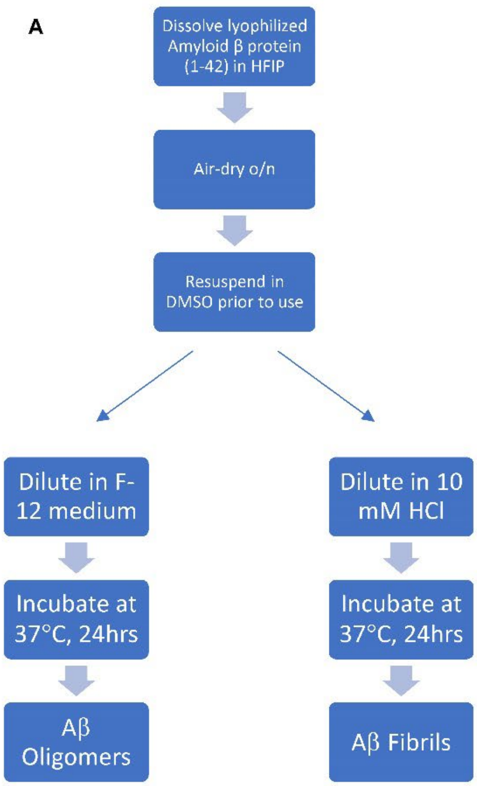

B

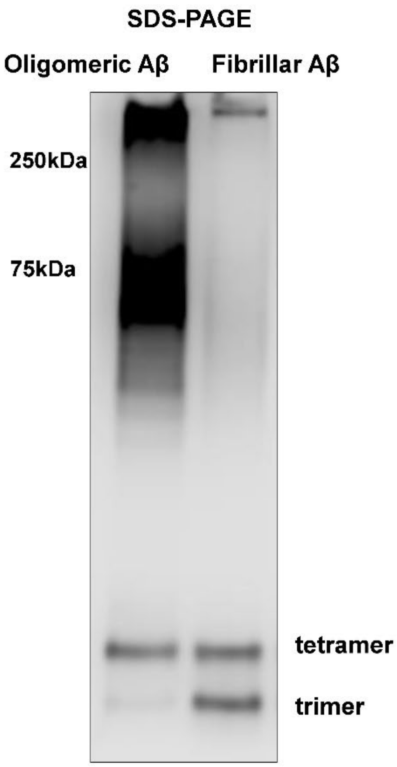

**Fig. S2. Generation of oligomeric and fibrillar Aβ.** (**A**) Schematic illustration of experimental procedure. (**B**) Oligomeric and fibrillar Aβ were confirmed by SDS-PAGE followed by probing with the anti-β-amyloid antibody clone 6E10.

### Supplementary Figure 3

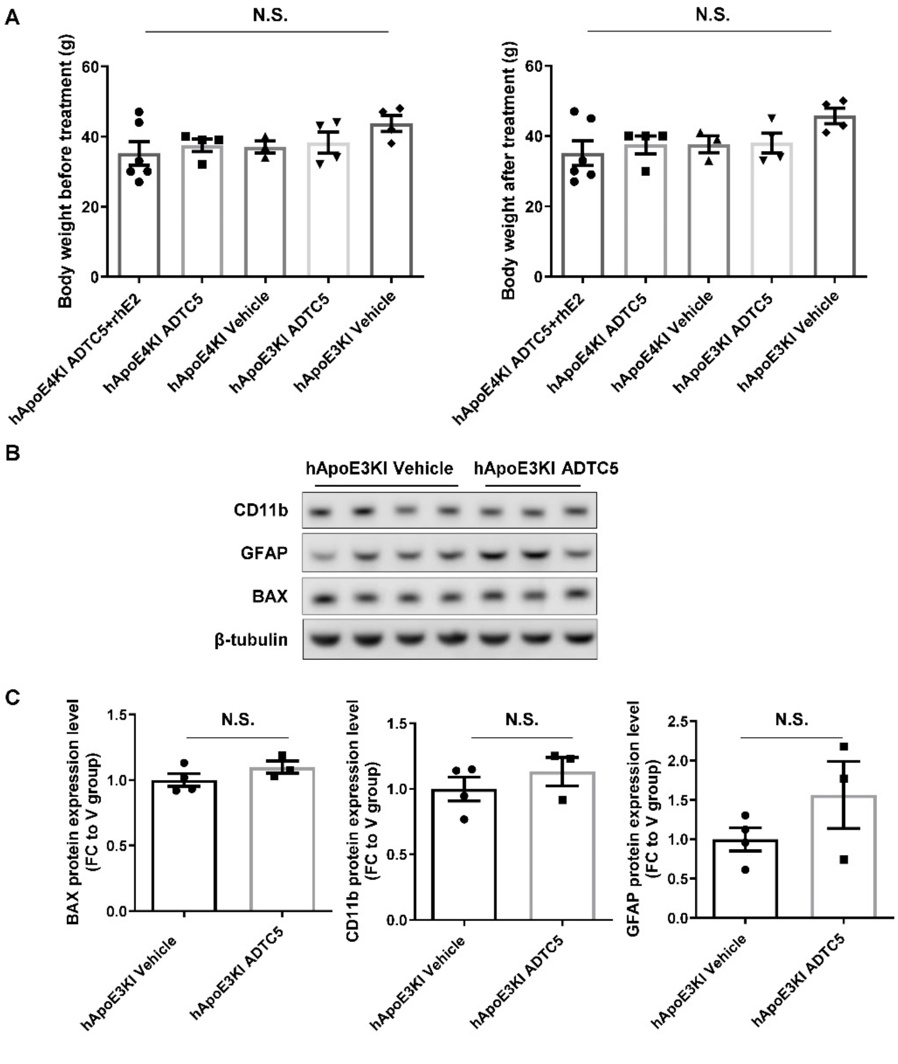

**Fig. S3. No significant changes in body weight or gliosis were observed in hApoE4KI and hApoE3KI mice after four weeks of weekly treatment with rhApoE2+ADTC5, ADTC5, or vehicle alone.** (**A**) Body weight records of all the groups before and after treatment. (**B-C**) GFAP, CD11b, and BAX expression levels in hApoE3KI mice treated with ADTC5 or vehicle alone. Data are presented as the group mean ± SEM. Group differences were analyzed using one-way ANOVA with Tukey's post-hoc test or Student's t-test. N.S., non-significant. N = 3-6 mice per group.

### Supplementary Figure 4

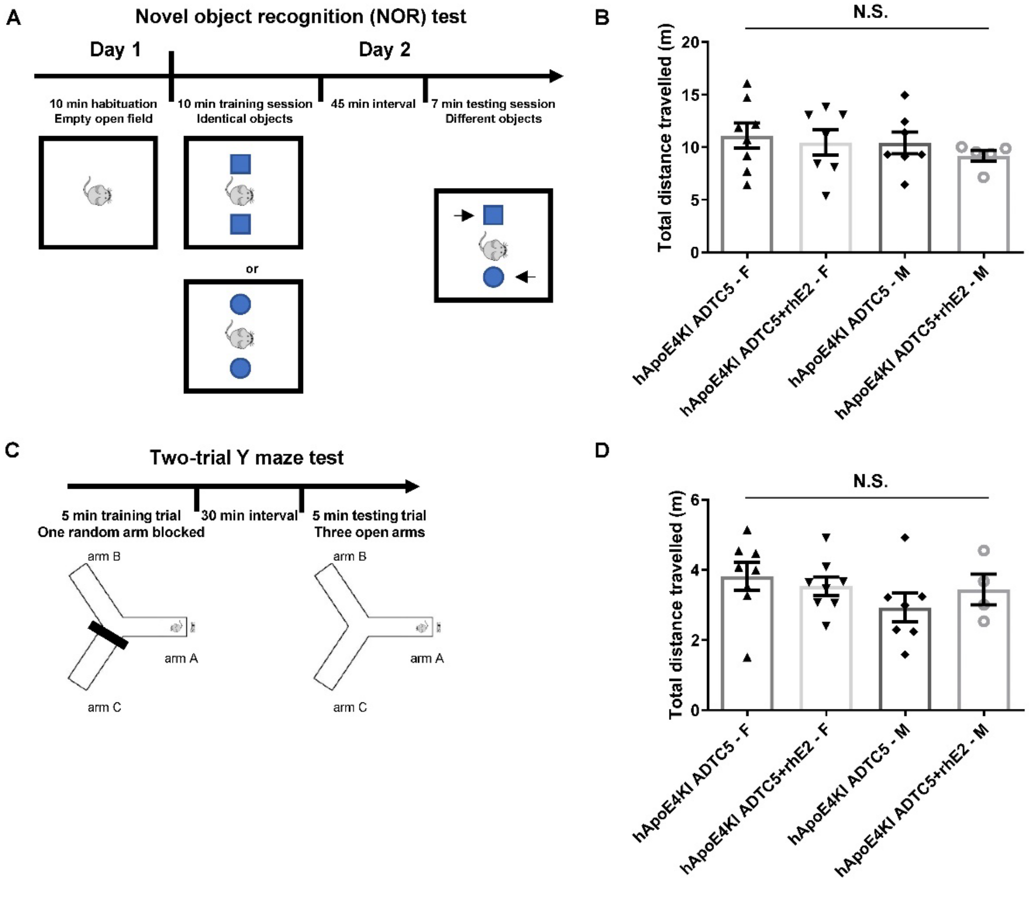

**Fig. S4. NOR and Y-maze two-trial behavioral tests were conducted on hApoE4KI mice after eight weeks of weekly treatment with rhApoE2+ADTC5 or ADTC5 alone.** (**A**) Schematic illustration of the NOR experimental design. (**B**) Total distance traveled in the open field did not differ among groups. (**C**) Schematic illustration of the Y-maze two-trial experimental design. (**D**) Total distance traveled in the Y-maze did not differ across groups. Data are presented as the group mean ± SEM. Group differences were analyzed using one-way ANOVA with Tukey's post-hoc test. N.S., non-significant. N = 4-8 mice per group.

### Supplementary Figure 5

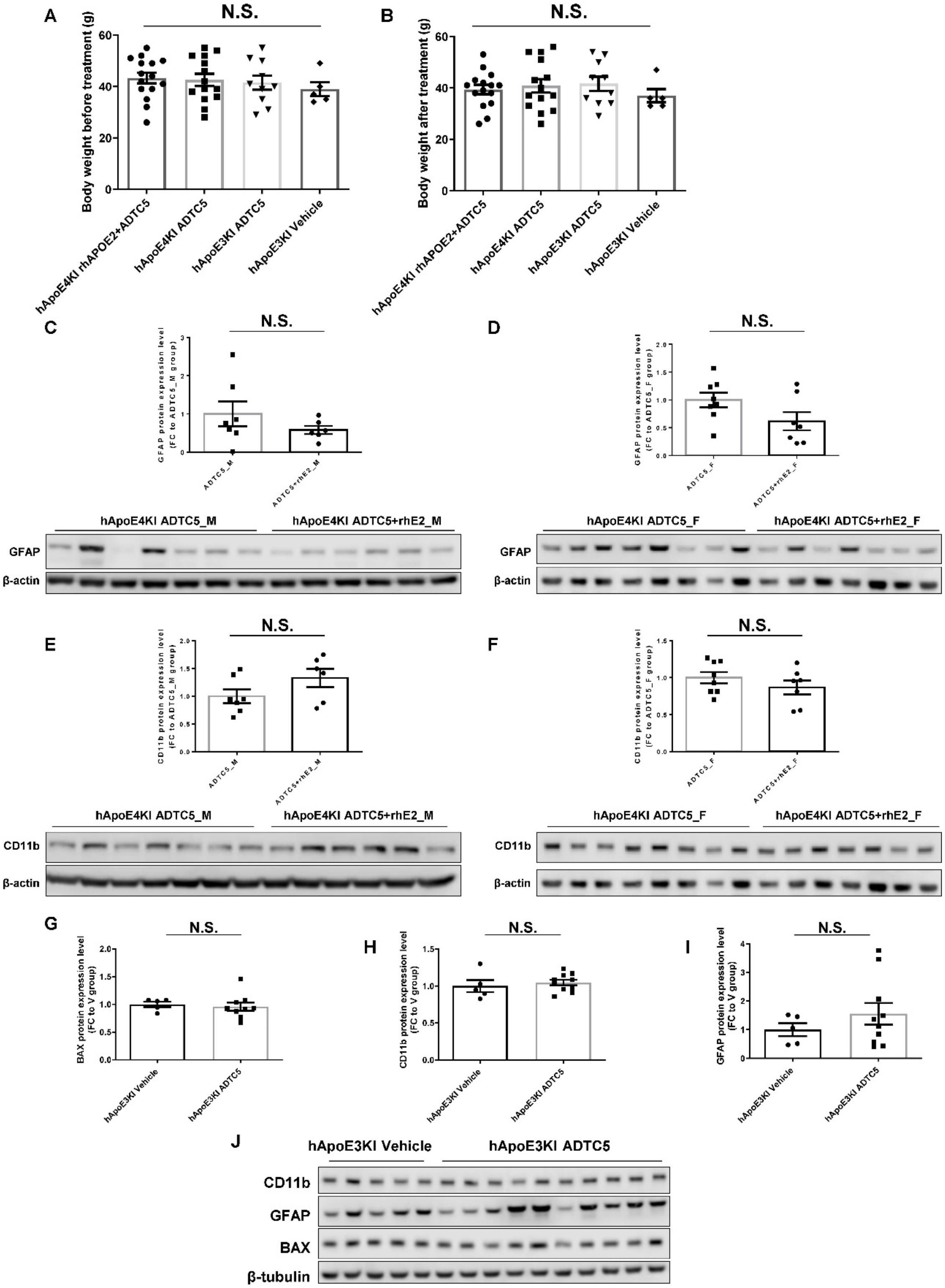

**Fig. S5. No significant changes in body weight or gliosis were observed in hApoE4KI and hApoE3KI mice after eight weeks of weekly treatment with rhApoE2+ADTC5, ADTC5, or vehicle alone.** (**A-B**) Body weight records for all the groups before and after treatment. (**C-F**) GFAP and CD11b expression levels in hApoE4KI mice treated with rhApoE2+ADTC5 or ADTC5 alone. (**G-J**) GFAP, CD11b, and BAX expression levels in hApoE3KI mice treated with ADTC5 or vehicle alone. N.S., non-significant. Data are presented as the group mean ± SEM. Group differences were analyzed using one-way ANOVA with Tukey's post-hoc test or Student's t-test. N = 5-15 mice per group.

**Supplementary Table 1**

| Lipid Class | Male rhApoE2+ADTC5 (G1) versus Male ADTC5 control (G2) | | | | |
|---|---|---|---|---|---|
| | FC (G1/G2) | T-Test (p) | VIP | FA1 standardized name | FA2 standardized name |
| ***Fatty acids*** | | | | | |
| FA(22:6) | 4.182386481 | 0.032276926 | 1.94684 | DHA | |
| ***Sphingolipids*** | | | | | |
| SPHP(m22:1) | 0.393575095 | 0.018791639 | 2.07789 | | |
| CerP(t44:7) | 0.360949101 | 0.017033915 | 2.09957 | | |
| SM(t42:3) | 2.672255874 | 0.04852048 | 1.83204 | | |
| ***Glycoglycerolipids*** | | | | | |
| MGDG(30:0) | 0.448732461 | 0.01934279 | 2.07159 | | |
| MGDG(39:6) | 0.462649404 | 0.026910173 | 1.99341 | | |
| ***Phospholipids*** | | | | | |
| PA(16:0_16:0) | 4.327321329 | 0.023912464 | 2.02231 | Palmitic acid | Palmitic acid |
| PC(18:1_20:4) | 2.291277398 | 0.026993007 | 1.99262 | Octadecenoic acid | Eicosatetraenoic acid |
| PC(20:0_18:1) | 2.734436347 | 0.035765283 | 1.91939 | Arachidic acid | Octadecenoic acid |
| PE(16:0_18:2) | 2.919261251 | 0.031953278 | 1.94953 | Palmitic acid | Octadecadienoic acid |
| PE(18:0_18:1) | 3.956700424 | 0.008382164 | 2.23735 | Stearic acid | Octadecenoic acid |
| PE(18:1_20:4) | 2.695900743 | 0.010917359 | 2.18939 | Octadecenoic acid | Eicosatetraenoic acid |
| PE(18:1e_22:4) | 3.616874912 | 0.010992313 | 2.18803 | - | Docosatetraenoic acid |
| PG(16:0_20:4) | 2.551597152 | 0.016198026 | 2.11041 | Palmitic acid | Eicosatetraenoic acid |
| PG(16:1_22:6) | 2.59411243 | 0.013417421 | 2.14929 | Hexadecenoic acid | DHA |
| PG(18:1_20:3) | 2.346761073 | 0.007031868 | 2.2673 | Octadecenoic acid | Eicosatrienoic acid |
| PG(34:1) | 2.419588863 | 0.007208595 | 2.26315 | Tetratriacontenoic acid | |
| PS(18:0_22:6) | 3.301719282 | 0.044025606 | 1.86083 | Stearic acid | DHA |
| ***Phospholipids*** | | | | | |
| PC(18:1_24:1) | 0.495967166 | 0.010860203 | 2.19036 | Octadecenoic acid | Tetracosenoic acid |
| PE(16:0_14:0) | 0.282478004 | 0.02966468 | 1.96885 | Palmitic acid | Myristic acid |
| PE(16:1e_22:6) | 0.10508052 | 0.044738328 | 1.85619 | - | DHA |
| PE(18:0_16:0) | 0.405423589 | 0.029881758 | 1.96704 | Stearic acid | Palmitic acid |
| PE(18:1_22:6) | 0.340856599 | 0.005784868 | 2.2988 | Octadecenoic acid | DHA |
| PE(18:1e_18:1) | 0.229901383 | 0.043303998 | 1.86559 | - | Octadecenoic acid |
| PE(18:2e_20:4) | 0.163364781 | 0.002991603 | 2.39299 | - | Eicosatetraenoic acid |
| PE(20:0_22:6) | 0.365349156 | 0.006293252 | 2.28538 | Arachidic acid | DHA |
| PE(22:6_22:6) | 0.434234478 | 0.023121255 | 2.03039 | DHA | DHA |
| PE(34:0) | 0.466489004 | 0.042137724 | 1.87359 | Gheddic acid | |
| PI(16:0_16:0) | 0.14786634 | 0.024178538 | 2.01972 | Palmitic acid | Palmitic acid |
| PI(18:4_22:4) | 0.491148423 | 0.037758198 | 1.90457 | Octadecatetraenoic acid | Docosatetraenoic acid |
| DLCL(38:5) | 0.060989642 | 0.016040579 | 2.11235 | | |

*FC: fold change; VIP: variable importance in projection; FA: fatty acid; SPHP: sphingosine phosphate; CerP: ceramide phosphate; SM: sphingomyelin; MGDG: monogalactosyldiacylglycerol; PA: phosphatidic acid; PC: phosphatidylcholine; PE: phosphatidylethanolamine; PG: phosphatidylglycerol; PI: phosphatidylinositol; PS: phosphatidylserine; DLCL: dilysocardiolipin*

**Supplementary Table 2**

| Lipid Class | Female rhApoE2+ADTC5 (G3) versus Female ADTC5 control (G4) | | | | |
|---|---|---|---|---|---|
| | FC (G3/G4) | T-Test (p) | VIP | FA1 standardized name | FA2 standardized name |
| ***Sphingolipids*** | | | | | |
| SPHP(m20:1) | 4.02124585 | 0.024873766 | 2.03946 | | |
| CerP(m38:1) | 2.070231518 | 0.010366135 | 2.22809 | | |
| Hex1Cer(d40:1) | 0.441181484 | 0.036451883 | 1.93939 | | |
| ***Phospholipids*** | | | | | |
| PA(16:0_16:0) | 3.121024348 | 0.021054784 | 2.07958 | Palmitic acid | Palmitic acid |
| PA(16:0_20:4) | 2.334768181 | 0.018031887 | 2.11466 | Palmitic acid | Eicosatetraenoic acid |
| PE(18:0_18:1) | 2.563683577 | 0.037683002 | 1.93046 | Stearic acid | Octadecenoic acid |
| PE(18:0_22:4) | 2.751627012 | 0.007186536 | 2.29371 | Stearic acid | Docosatetraenoic acid |
| PE(18:0_22:5) | 3.26905677 | 0.00392282 | 2.38763 | Stearic acid | Docosapentaenoic acid |
| PE(18:0_22:6) | 2.630864091 | 0.023752845 | 2.05084 | Stearic acid | DHA |
| PE(18:2e_20:1) | 2.907025628 | 0.013598258 | 2.17506 | - | Eicosenoic acid |
| PE(32:1_22:6) | 2.203347664 | 0.001437027 | 2.51211 | Dotriacontenoic acid | DHA |
| PEt(34:1) | 2.018074885 | 0.005232733 | 2.34495 | Tetratriacontenoic acid | |
| PEt(38:6) | 2.094326549 | 0.005210198 | 2.3454 | Octatriacontahexaenoic acid | |
| PS(18:0_20:2) | 2.253741064 | 0.041881731 | 1.89997 | Stearic acid | Eicosadienoic acid |
| PS(41:6) | 2.061696962 | 0.0491097 | 1.85244 | - | |
| ***Phospholipids*** | | | | | |
| PA(18:0_22:6) | 0.468775388 | 0.031592451 | 1.97841 | Stearic acid | DHA |
| PA(20:4_22:6) | 0.261441935 | 0.015283783 | 2.15048 | Eicosatetraenoic acid | DHA |
| PE(16:1e_18:1) | 0.421914232 | 0.013397539 | 2.17789 | - | Octadecenoic acid |
| PE(18:0_16:0) | 0.445749325 | 0.008642639 | 2.26156 | Stearic acid | Palmitic acid |
| PE(18:1_18:1) | 0.407152711 | 0.042663205 | 1.89464 | Octadecenoic acid | Octadecenoic acid |
| PE(18:1e_22:6) | 0.469145712 | 0.037962993 | 1.92834 | - | DHA |
| PE(18:2e_18:1) | 0.338696366 | 0.036250936 | 1.94103 | - | Octadecenoic acid |
| PE(20:0e) | 0.306620713 | 0.030625774 | 1.98664 | - | |
| PE(20:1e) | 0.465876132 | 0.045229134 | 1.87747 | - | |
| PG(34:1) | 0.306405977 | 0.044656109 | 1.88143 | Tetratriacontenoic acid | |
| PS(18:0_18:1) | 0.375133368 | 0.016944253 | 2.12855 | Stearic acid | Octadecenoic acid |
| PS(40:6) | 0.482491013 | 0.018095788 | 2.1141 | | |

*FC: fold change; VIP: variable importance in projection; SPHP: sphingosine phosphate; CerP: ceramide phosphate; Hex1Cer: hexosylceramide; PA: phosphatidic acid; PE: phosphatidylethanolamine; PG: phosphatidylglycerol; PS: phosphatidylserine*
